# An expanded metabolic pathway for androgen production by host-associated bacteria

**DOI:** 10.1101/2024.06.09.598130

**Authors:** Taojun Wang, Saeed Ahmad, Angélica Cruz-Lebrón, Sarah E. Ernst, Kelly Yovani Olivos Caicedo, Yoon Jeong, Briawna Binion, Pauline Mbuvi, Debapriya Dutta, Francelys V. Fernandez-Materan, Adam M. Breister, Jae Won Lee, Jason D. Kang, Spencer C. Harris, Shigeo Ikegawa, H. Rex Gaskins, John W. Erdman, Glen Yang, Isaac Cann, Steven L. Daniel, Phillip B. Hylemon, Karthik Anantharaman, Rafael C. Bernardi, João M.P. Alves, Karen S. Sfanos, Joseph Irudayaraj, Jason M. Ridlon

## Abstract

A growing body of literature implicates host-associated microbiota in the modulation of circulating androgen levels in the host, which could have far-reaching implications for androgen-mediated diseases. However, the microbial genetic pathways involved in androgen production remain unknown. Here, we report the first host-associated microbial gene (*desF*) encoding an enzyme that catalyzes conversion of androstenedione to epitestosterone (epiT) in the gut bacterium, *Clostridium scindens*. Despite current dogma that epiT is a nuclear androgen-receptor (AR) antagonist, we demonstrate that epiT is a potent androgen, as assessed by its ability to promote prostate cancer cell growth and expression of prostate specific antigen (PSA). We then quantified the *desF* gene in fecal samples collected from individuals with advanced prostate cancer (rising blood PSA) undergoing androgen deprivation therapy combined with abiraterone acetate and prednisone (AA/P). Strikingly, fecal *desF* levels were elevated in a subset of individuals progressing on AA/P versus samples taken during AA/P response (stable). Importantly, we observed that AA does not inhibit the bacterial desmolase enzyme that is analogous to the human drug target of AA. We then determined that bacterial isolates from urine or prostatectomy tissue are capable of androgen production. From these isolates we detected 17β-hydroxysteroid dehydrogenase (17β-HSDH) activity, which has not been previously reported in urinary tract bacteria, and discovered the *desG* gene in urinary isolates encoding 17β-HSDH that catalyzed conversion of androstenedione to testosterone. Applying advanced artificial intelligence and molecular dynamics, we predict the structures and ligand binding to DesF and DesG. Using a novel bioengineered microencapsulation technique, we demonstrate that urinary androgen-producing bacterial strains can also promote prostate cancer cell growth through steroid metabolism. Taken together, our results are a significant advance for steroid microbiology in humans and suggest that these microbial biotransformations should be further studied in the context of androgen-mediated physiological processes and diseases.

## Introduction

Human microbiome research has revealed that the human body is composed numerically of equal bacterial and mammalian cells^1^, and that 99% of the functional genes are microbial^2^. Accordingly, the functional capacity of the human-associated microbiota to influence human health and disease is immense. For example, a growing body of literature implicates the gut microbiota in the modulation of circulating androgen levels in the host^3,4^, which could have far-reaching implications for androgen-mediated diseases, as highlighted in studies using murine models. The nonobese diabetic (NOD) mouse model recapitulates spontaneous autoimmunity-mediated type 1 diabetes. Fecal microbiota transplant (FMT) of gut microbiota from adult male to immature female NOD mice altered the recipient’s microbiota, resulting in elevated circulating testosterone (T), metabolomic changes, reduced islet inflammation, and lowered autoantibody production, resulting in robust type 1 diabetes protection^3^. These effects were driven by gut microbiota-mediated T production. Another study reported that FMT from metastatic castration resistant prostate cancer (mCRPC) donors promotes castration-resistant tumor growth in mice^5^. Antibiotic treatment likewise delayed the onset of castration resistance in both immune-competent and immune-deficient mouse models. Results from both an in vitro culture system and in vivo androgen-tracing experiments implicated the gut microbiota as an important source of androgens including T and dehydroepiandrosterone (DHEA); however, the bacterial genes encoding the enzymes involved in this androgen production were not determined^5^. Studies such as these establish that host-associated microbial communities are significant contributors to circulating androgens, and that gut androgen production may influence androgen-mediated disease. Importantly, the bacterial enzymatic pathways that mediate gut androgen production are not fully elucidated, and the characterization of these pathways offers novel approaches for therapeutic targeting.

Using transcriptomics, we discovered the first gut bacterial enzymatic pathway (*desABC* operon) that is involved in the conversion of cortisol derivatives to 11-oxy-androgens in *Clostridium scindens*^6,7^. We demonstrated that *desC* encodes steroid 20ɑ-HSDH involved in side-chain oxidoreduction^6,8^, while *desAB* encodes steroid-17,20-desmolase involved in side-chain cleavage of cortisol derivatives to 11-oxy-androgens^6,7^. 11-oxy-androgens are now regarded as potent androgen receptor (AR) ligands (e.g., 11-keto-T and 11-keto-dihydrotestosterone (11-keto-DHT) on par with T and DHT^9^. We theorize that androgen production in the gut via bacterial species that carry the *des* operon may contribute to disease etiology, with prostate cancer as one exemplary paradigm. mCRPC is largely incurable and characterized by progressive metastatic cancer growth despite treatment that blocks androgen synthesis (e.g., gonadotropin-releasing hormone agonist/antagonist) in the testes. Second-line treatments that block adrenal androgen synthesis (e.g., abiraterone acetate (AA) given with the replacement glucocorticoid prednisone (P)), and/or directly antagonize AR (e.g., enzalutamide) are likewise not curative, and resistance invariably develops^10^. Despite castrate levels of circulating T in individuals with mCRPC, intra-tumoral levels of androgens remain high^11, 12^. Current research on the source(s) of these intra-tumoral androgens is almost entirely focused on host enzymatic biosynthesis and intracrine pathways through which androgen-precursors are synthesized and become AR-ligands^13–16^. Despite these efforts, the source of intra-tumoral androgens that may counteract therapeutic castration is unclear. Furthermore, the capacity for androgen production by microbiota colonizing other host epithelial sites to contribute to disease etiology and therapeutic response is completely unexplored.

Herein, we report an expansion of the bacterial desmolase pathway in *desAB* harboring taxa to include host-associated bacteria expressing 17ɑ-hydroxysteroid dehydrogenase (17ɑ-HSDH; *desF*) or 17β-HSDH (*desG*) involved in the formation of epitestosterone (epiT) and T, respectively. We also demonstrate that, contrary to the current dogma, epiT is a potent androgen that drives prolonged expression of kallikrein-3 (KLK3, also known as prostate specific antigen or PSA) and proliferation of prostate cancer cell lines. In the clinical setting, we found that fecal *desF*-carrying bacteria are present in the gut microbiota of individuals with advanced prostate cancer, and that fecal *desF* levels are elevated in individuals with disease progression on androgen-deprivation therapy combined with AA/P versus those who are responding. Since we previously demonstrated that the genome of urinary strains of *Propionimicrobium lymphophilum* can harbor DesAB^17^, we isolated strains of *P. lymphophilum* from human prostatectomy tissue and urine and found them to be capable of converting cortisol and prednisone to AR ligands that drive hormone-responsive prostate cancer cell (LNCaP) proliferation *in vitro* using a unique microencapsulation technique. These findings significantly expand our knowledge of human steroid microbiology and provide clinically relevant evidence that this expanded steroid-17,20-desmolase pathway should be further investigated.

## Results

### A novel pathway for the formation of epiT by the gut microbiota

*C. scindens* ATCC 35704 (*Csci*35704) expresses steroid-17,20-desmolase encoded by the *desAB* genes^6,7^. Three decades ago *C. scindens* VPI 12708 (*Csci*12708) was reported to convert androstenedione (AD) to epiT indicating the presence of a gene encoding the enzyme 17ɑ-hydroxysteroid dehydrogenase (17ɑ-HSDH)^18^. We reasoned that co-culture of both strains in the presence of 11-deoxycortisol (11DC) would yield epiT with a stable AD intermediate. As predicted, co-culture of *Csci12708* and *Csci35704* yielded the expected conversion of 11DC (RT 3.81 min; 347.2 *m/z*) to both AD (RT 4.47 min; 287.2 *m/z*) and epiT (RT 4.60; 289.2 *m/z*) after 24 h (**Fig.1a, b**). 11DC (0 h = 45.54 ± 0.59 µM) was depleted in 24 h, yielding AD (24 h = 6.20 ± 0.22 µM) and epiT (24 h = 27.42 ± 0.64 µM). We confirmed the formation of epiT from AD in pure cultures of *Csci*12708 by a combination of high-resolution LC/MS/MS, and proton and carbon NMR (**Extended Data Fig. 1**). Endocrine pathways in steroidogenic tissues (e.g., adrenal gland and gonads) generate AD and epiT through pathways distinct from *Csci* strains. Indeed, DHEA is converted to AD via HSD3B2 or the lyase activity of CYP17A1 coupled with P450 oxidoreductase (POR) and cytochrome b5 (CYB5) generates AD from 17ɑ-hydroxyprogesterone^19^. While the metabolic pathway for epiT formation by human steroidogenic pathways is unknown, it is speculated to derive from 5-androstene-3β,17ɑ-diol rather than AD^20^.

**Figure 1.**
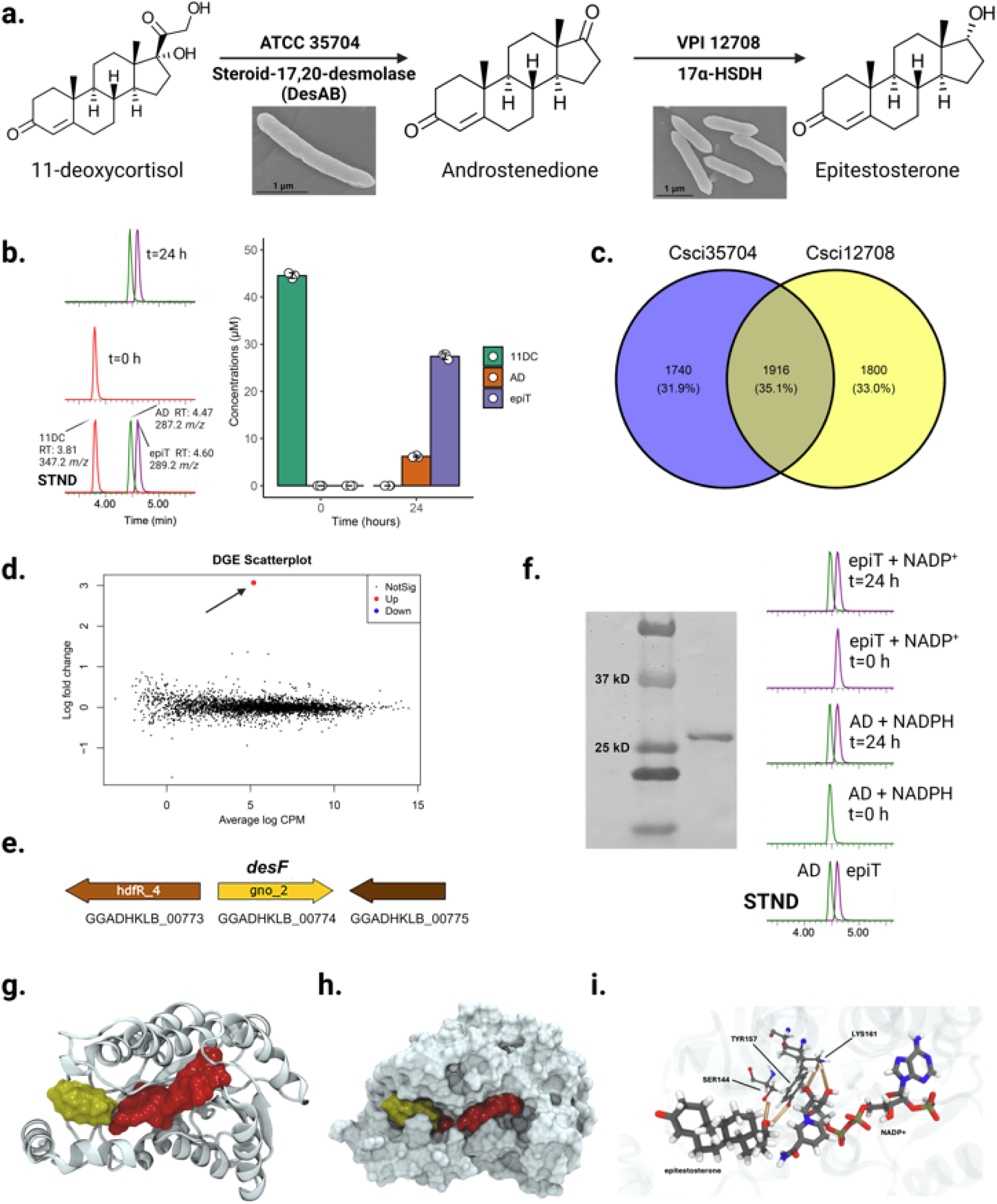
Identification of the *desF* gene encoding steroid 17ɑ-hydroxysteroid dehydrogenase involved in epitestosterone formation by *Clostridium scindens*. a. proposed biochemical pathway by which *Csci*35704 and *Csci*12708 convert 11DC to AD and epiT. SEM images of *Csci*35704 and *Csci*12708 are included. **b.** LC/MS/MS chromatographs of 11DC, AD, and epiT at time 0 and 24 h (left) and quantification of metabolites (right) (n=3). **c.** Venn diagram summarizing comparative genomic analysis between *Csci*35704 and *Csci*12708 (see Extended Data Fig. 2 for reductases unique to *Csci*12708). **d.** Scatterplot of RNA-Seq analysis to identify genes upregulated by 50 µM androstenedione in *Csci*12708. Significantly upregulated genes (>0.58 log_2_fold; <0.05 FDR) in red, downregulated genes in blue, and not differentially regulated in black. **e.** Gene organization of 17ɑ-hydroxysteroid dehydrogenase candidate (*“desF*”) identified in RNA-Seq analysis. **f.** SDS-PAGE of Streptavidin-purified recombinant DesF and LC/MS/MS chromatographs showing NADPH-dependent conversion of AD to epiT at 0 and 24 h, and NADP^+^-dependent conversion of epiT to AD at 0 h and 24 h. **g.** Ribbon diagram of AlphaFold 2 structural prediction of DesF. NADP^+^ (red) and epitestosterone (yellow) are depicted as space-filling models, and were aligned and placed with VMD and ligand-structure interactions were minimized using NAMD through its QuikMD interface. **h.** Space-filling model of both the DesF (gray) and ligands NADP^+^ (red) and epitestosterone (yellow). **i.** Molecular dynamic trajectory analysis revealed strong interactions between ligands and catalytic triad. SER145 and TYR160 form stable hydrogen bonds with epitestosterone, while LYS164 was identified as key in NADP^+^ stabilization.

### The gut microbial desF gene encodes a novel 17ɑ-HSDH

We next sought to identify the gene(s) encoding 17ɑ-HSDH in *Csi12708* responsible for catalyzing the conversion of AD to epiT. We performed comparative genomics between *Csci*35704 and *Csci*12708 to identify reductases unique to *Csci*12708 (**Fig. 1c; Extended Data Fig. 2).** The strains share 35% of their genes (1916 ORFs) with 33% of genes (1,800 ORFs) unique to *Csci*12708 (**Fig. 1c**). We narrowed this list down to three protein families known to include HSDH enzymes: 25 belonging to the short chain dehydrogenase/reductase (SDR) family; 23 to the medium chain dehydrogenase/reductase (MDR) family; and 2 to the aldo-keto reductase (AKR) family^20^. Of these, 18 SDR, 18 MDR, and 2 AKR proteins are unique to *Csci*12708 (**Extended Data Fig. 2**).

Given this relatively large number of candidates and the reported steroid-inducibility of 17ɑ-HSDH^18^, we opted to utilize genome-wide transcriptomics to identify candidates after growth of *Csci*12708 in the presence of 50 µM 11β-hydroxyandrostenedione (11OHAD) (n=4) vs. uninduced controls (n=4) (**Fig. 1d**). The addition of 11OHAD significantly upregulated (3.07 log_2_FC; FDR 0.012) the expression of a single gene (GGADHKLB_00774) (**Fig. 1d**). This observation was reproducible since repeat transcriptome analysis with additional biological replicates (n=3) yielded the same overall result (**Extended Data Fig. 3; Supplementary Table 1**). GGADHKLB_00774 is predicted to encode a 27-kDa short chain dehydrogenase/reductase (SDR) superfamily protein (**Fig. 1e**). Importantly, GGADHKLB_00774 is in the list of reductases found in *Csci*12708 but not in *Csci*35704 (**Extended Data Fig. 2).** Since many HSDHs are represented in the SDR superfamily^21^, we cloned GGADHKLB_00774 in a pET51b(+) vector for overexpression of the recombinant streptavidin-tagged enzyme. With 10 nM purified rGGADHKLB_00774 (30.105 kDa determined by proteomics), AD was converted to epiT in the presence of NADPH (but not NADH) (**Fig. 1f**), establishing that GGADHKLB_00774 from *Csci12708* encodes a novel NADPH-dependent 17ɑ-HSDH in the SDR superfamily. We propose the name *desF* for this gene, which constitutes the first host-associated microbial gene encoding a steroid 17ɑ-HSDH.

To investigate the predicted molecular mechanism of ligand binding in the enzyme DesF we utilized AlphaFold 2^22^ through its QwikFold interface in VMD^23^. Using VMD, the structure predicted by AlphaFold were aligned with similar structures available in the Protein Data Bank (PDB). Following a previously established protocol^8^, we used homologous structures from the PDB to fit an NADP^+^ molecule into the binding pockets of DesF monomer. We obtained homologous structures (PDB IDs: 4ILK, 4EJ6, 4A2C, 3QE3, 3GFB, 2DQ4, 2DFV, 2D8A, 1PL7, and 1E3J) using protein BLAST. The alignment and placement of the NADP^+^ molecule on the binding site was performed using VMD. Utilizing advanced options in QwikMD^24^, the ligand structure was minimized in the pocket along with nearby protein residues, while most of the protein structure remained static.

The structure of epiT in DesF was also fitted to the most probable binding site using VMD. Docking was performed using a combined manual and computational approach, where VMD was used to position reference atoms, and NAMD^25^ through its QwikMD interface was used to minimize the structure of the complex, using a previously established protocol^8^. The complex was then solvated and subjected to a 100 ns equilibrium MD simulation. AI-based predictions and MD simulations showed that both epiT (yellow) and NADP^+^ (red) occupy a mostly open cleft (**Fig. 1g, h**).

MD simulations can be used to investigate the stability of molecular systems and predict the behavior and function of proteins or protein domains. Here, after 100 ns of MD simulation, we observed that epiT, and NADP^+^ remained stable in the predicted catalytic cleft. Analysis of the MD trajectory revealed strong interactions between the ligands and serine, tyrosine, and lysine residues in DesF. SER145 and TYR160 formed stable interactions with the epiT molecule (**Fig. 1i**). Additionally, LYS164 was identified by our network analysis as a key residue in stabilizing NADP^+^ in its position.

Phylogenetic analysis indicates that DesF appears to be unique to *C. scindens* strains, with these strains clustering in a clade well separated from the other sequences analyzed (**Extended Data Fig. 4**). Accordingly, amino acid identity drops precipitously to at most ∼50% identity in SDR family proteins in other taxa based on the BLAST results in NCBI. DesF shares only 20.2% amino acid identity with *Mus musculus* 17ɑ-HSDH, which is in the aldo-keto reductase family (AKR)^26^. We thus sought to determine the proportion of *C. scindens* strains that harbor *desABC* and *desF* genes. We sequenced the genomes of 14 *C. scindens* strains and obtained an additional 20 genomes from sequenced *C. scindens* strains from NCBI^27–29^. Interestingly, two clades of *C. scindens* strains were apparent in whole genome phylogeny (**Extended Data Fig. 5**) with the two most studied strains, *Csci*35704 (Clade 1) and *Csci*12708 (Clade 2), being distinct. In Clade 1, 12 strains harbored *desABC* genes, with *desF* being absent. In Clade 2, 17 of 19 strains encode *desF*, and two strains, SL.1.22 and S076 had both *desABC* and *desF* genes.

We then constructed metagenome assembled genomes (MAGs) from publicly available human metagenomes resulting in 225 *C. scindens* MAGs (**Supplementary Table 2**). In the *C. scindens* MAGs, 20 had *desABC*, 97 had *desF*, and 4 had both *desABC* and *desF*(**Supplementary Table 2**). This indicates that strain variation in the gut microbiomes of humans may identify a subset of individuals with higher potential to convert glucocorticoids into androgens, including derivatives of epiT.

### EpiT serves as an AR agonist that promotes prostate cancer cell proliferation

EpiT is regarded as an “antiandrogen” that is expected to bind to and antagonize AR and reduce prostate cancer cell growth^30^. This dogma has been challenged with a recent study indicating that epiT instead serves as an AR agonist in a reporter cell line^31^. Circulating epiT is measured in the low nanomolar concentrations, with epiT/T ratios of 0.1 for women and 1 in men^32^. However, little evidence in the literature has examined epiT for its potential to alter cell physiology via nuclear AR^30^. We thus compared the 96-h growth of androgen-sensitive prostate cancer cells (LNCaP) grown in charcoal-stripped medium in the presence of either 1 nM or 10 nM AD, T, and epiT to a vehicle control (VC; 0.5% v/v methanol) (**Fig. 2a**). As expected, at 1 nM, T caused significant proliferation (1.46 ± 0.11 fold; *P* = 2.0 × 10^-07^) relative to VC (n= 6); while the androgen-precursor, and non-AR ligand, AD, did not (0.90 ± 0.14 fold; *P* = 0.123) (**Fig. 2b**). Unexpectedly, EpiT at 1 nM also caused significant proliferation relative to both VC (1.76 ± 0.079 fold; *P* = 1.1 × 10^-10^) and T (*P* = 8.2 × 10^-05^) (**Fig. 2b**). A dose-dependent effect was also observed such that at 10 nM, AD caused significant proliferation relative to VC (1.29 fold ± 0.23 fold; *P* = 0.00523) (**Fig. 2b**). Proliferation caused by T at 10 nM increased significantly with respect to T at 1 nM (*P* = 0.0024), and proliferation was significant with respect to both AD (*P* = 3.8 × 10^-05^) and VC (*P* = 7.8 × 10^-08^) (**Fig. 2b**). This pattern continued with epiT treatment at 10 nM in that proliferation was significant relative to 1 nM epiT (*P* = 2.0 × 10^-05^). EpiT treatment at 10 nM caused proliferation significantly above all other treatment groups (VC, *P* = 1.4 × 10^-10^; AD, *P* = 7.8 × 10^-09^; T, *P* = 0.00018) (**Fig. 2b**).

**Figure 2.**
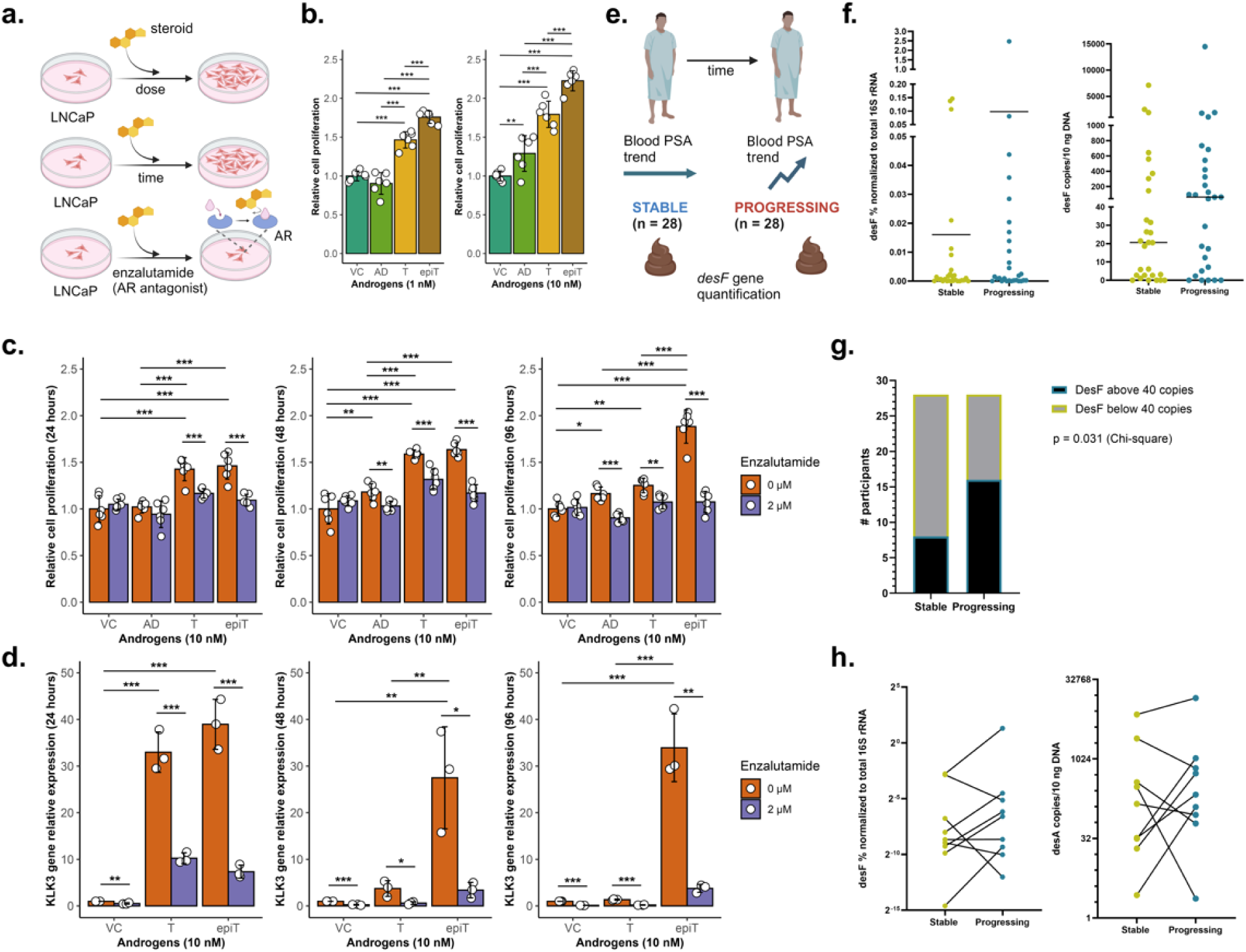
Epitestosterone and gut microbial *desF* in prostate cancer. **a.** Schematic representation of in vitro experiments examining dose and time dependent growth of LNCaP cells in the presence of androgen candidates as well as determination of the effect of androgen receptor antagonist, enzalutamide. **b.** Dose-dependent proliferation (MTS assay) of LNCaP cells in the presence of androstenedione (AD), testosterone (T), epitestosterone (epiT), and vehicle control (VC; methanol 0.5% v/v) at 1.0 nM or 10.0 nM (n = 6). **c.** Time-dependent proliferation (MTS assay) of LNCaP cells in the presence (2 µM) or absence of AR-antagonist, enzalutamide. Proliferation was measured at 24, 48, or 96 h in the presence of 10 nM steroids or VC (n = 6). **d.** qRT-PCR quantification of *KLK3* gene (encoding prostate specific antigen, PSA) after treatment with 10 nM T, epiT, or VC at 24, 48, 96 h in the presence of absence of 2 µM enzalutamide (n=3). **e.** Schematic representation of clinical study examining fecal *desF* quantification (qPCR) in advanced prostate cancer patients undergoing treatment with AA/P either stable (blood PSA trend stable) (n = 28) or progressing (blood PSA trend increasing) (n = 28). **f.** Scatterplots of *desF* % normalized to total fecal 16S rRNA gene (left) or *desF* copy number/10 ng DNA in stable vs. progressing male patients. **g.** Proportion of stable vs. progressing patients above a threshold of 40 *desF* copies vs. below 40 *desF* copies. **h.** Fecal *desF* quantification (n = 12) of donor samples taken both while they were stable on AA/P and when they were progressing. In vitro data are shown with mean ± S.D. from six biological replicates. *P* values were calculated by unpaired *t*-test and Benjamini-Hochberg correction, * *P <* 0.05, ** *P <* 0.01,*** *P* < 0.001, **** *P* < 0.0001. Clinical data significance was determined by Chi squared analysis.

We next examined LNCaP proliferation over three time points. At 24 and 48 h, both T and epiT caused significant proliferation relative to AD (T, 24 h, *P* = 1.9 × 10^-05^; epiT, 24 h, *P* = 1.3 × 10^-05^; T, 48 h, *P* = 1.6 × 10^-06^; epiT, 48 h, *P* = 4.2 × 10^-07^) and VC (T, 24 h, *P* = 1.3 × 10^-05^; epiT, 24 h, *P* = 1.3 × 10^-05^; T, 48 h, *P* = 1.0 × 10^-08^; epiT, 48 h, *P* = 5.4 × 10^-09^) but were not significantly different in their effects relative to each other (**Fig. 2c**). At 96 h, T-stimulated proliferation decreased, although proliferation remains significant relative to VC (1.25 fold ± 0.078; *P* = 0.0012); however, proliferation caused by epiT at 96 h increases significantly relative to 24 and 48 h, and proliferation in the presence of epiT is highly significant relative to all other treatments (VC, *P* = 6.7 × 10^-11^; AD, *P* = 1.3 × 10^-09^; T, *P* = 8.0 × 10^-09^) (**Fig. 2c**). Since LNCaP cells express a mutant androgen receptor with broadened steroid-binding capacity^33^, we repeated these experiments with androgen-responsive VCaP cells that express wild type AR^33^. Significant proliferation (AD, 2 d, *P* = 0.00035; T, 2 d, *P* = 8.5 × 10^-07^; epiT, 2 d, *P* = 1.9 × 10^-8^; AD, 4 d, *P* = 1.4 × 10^-10^; T, 4 d, *P* = 3.6 × 10^-11^; epiT, 4 d, *P* = 3.6 × 10^-11^; AD, 8 d, *P* < 2.0 × 10^-16^; T, 8 d, *P* < 2.0 × 10^-16^; epiT, 8 d, *P* = 9.6 × 10^-14^) was observed with 10 nM AD, T, and epiT after 2, 4 and 8 days of exposure (**Extended Data Fig. 6**). These results demonstrate that epiT functions as an AR agonist that drives the proliferation of prostate cancer cells.

To determine whether epiT-induced proliferation requires AR agonism, we treated LNCaP cells with 2.0 µM enzalutamide, an AR competitive inhibitor and prostate cancer drug (IC_50_ 21.4 nM for LNCaP cells)^34^. Treatment with enzalutamide caused consistent and significant growth inhibition of LNCaP cells in the presence of 10 nM AD, T, or epiT at all time points, indicating that proliferation in all cases was AR-dependent (24 h, T, *P* = 0.00078, epiT *P* = 0.00017; 48 h, AD, *P* = 0.0053, T, *P* = 0.00025, epiT *P* = 1.7 × 10^-06^; 96 h, AD, *P* = 2.1 × 10^-05^, T, *P* = 0.0022, epiT *P* = 2.9 × 10^-06^) (**Fig. 2c**). We confirmed AR-dependent gene expression through measurement of the AR downstream gene target prostate specific antigen (PSA)/kallikrein-3 (KLK3)^35^. PSA expression at 96 h was increased to 33.92 ± 7.26-fold by epiT but reduced to 3.77 ± 0.86-fold by enzalutamide (**Fig. 2d**). PSA gene expression is elevated to 32.97 ± 4.33-fold in the presence of T at 24 h but dropped significantly to 3.74 ± 1.70-fold at 48 h and to 1.35 ± 0.12-fold by 96 h. In contrast to T, epiT caused prolonged PSA gene expression throughout the time course. In all cases, PSA gene expression is significantly reduced by enzalutamide treatment (**Fig. 2d**). Taken together, our results strongly indicate that epiT has a potent androgenic function not recognized previously.

### Measurement of human fecal desF gene indicates the potential physiological importance of gut bacterial 17ɑ-HSDH

We previously reported that prednisone (a replacement glucocorticoid commonly given in combination with AA) can be converted to 1,4-androstadiene-3,11,17-trione (AT) by DesAB^17^. We now questioned whether DesF could convert AT to epiAT, and whether treatment with this compound would lead to prostate cancer cell proliferation (**Extended Data Fig. 7**). We cultivated *Csci*12708 in the presence or absence of 50 µM AT, diluted the 0.2 µm-filtered, 72-h spent culture medium in sterile charcoal stripped RPMI medium, and added it to LNCaP cell culture at final concentrations of 0.1, 1.0, or 10.0 nM (**Extended Data Fig. 7a**). We verified quantitative depletion of AT after 48 h of cultivation with *Csci12708* (**Extended Data Fig. 7b, c**). Compared to spent medium (no steroid added) and spent medium with added AT (spiked in control), spent medium in which *Csci*12708 converted AT to epiAT resulted in dose-dependent proliferation of LNCaP cells, which was significant at 10 nM (CL, 1.66 ± 0.14 fold, *P* = 1.4 × 10^-06^) (**Extended Data Fig. 7d**). This proliferation was ablated by enzalutamide treatment indicating that epiAT triggered proliferation through AR-dependent signaling (**Extended Data Fig. 7e**).

We next determined whether bacteria carrying the *desF* gene are present in the gut microbiota of men being treated for advanced prostate cancer with AA/P, and whether quantitatively, fecal *desF* levels correlate with response to AA/P **(Supplementary Table 3**). Quantitative PCR (qPCR) primers were designed to target the *desF* gene, and we performed qPCR analysis on 56 fecal samples collected from 44 individuals with advanced prostate cancer undergoing treatment with AA/P (**Fig. 2e; Supplementary Table 3**). We found that 84.1% of the donors (37/44) had detectable fecal *desF*. Although not statistically significant, both the percentage of *desF* detected normalized to total 16S rRNA (as a surrogate for total bacterial load) as well as the absolute copy number of fecal *desF* were elevated in samples taken while individuals were progressing on AA/P versus samples taken during AA/P response (stable) (**Fig. 2f**). We observed that the majority of the samples in the AA/P stable group were quantitatively below 40 copies of *desF*. When the samples were categorized as below 40 copies versus above, the AA/P progressing group had significantly more samples with >40 copies of the *desF* gene detected (Chi-square, *P = 0.031*, **Fig 2g**). Twelve of the donors had samples taken both while they were stable on AA/P and when their disease was progressing. We found that a subset of these individuals had substantial increases in fecal *desF* levels during PSA progression on AA/P (**Fig. 2h**). These results suggest that *desF*-mediated androgen production by the gut microbiota (e.g., epiT) and/or metabolism of a replacement glucocorticoid (e.g., P) given in combination with AA could plausibly influence treatment response to AA/P, at least in a subset of patients. Thus, our discovery of *desF* and development of *desF* qPCR primers pave the way for future studies to examine bacterial epiT production as it relates to human health and disease.

### Formation of T derivatives from glucocorticoids by a bacterium isolated from prostatectomy tissue

We previously determined that *P. lymphophilum*, a normal inhabitant of the urinary tract, can harbor *desAB* genes^17^. We therefore sought to further explore microbial steroid biotransformations by urinary tract isolates. After screening pure cultures established in a prior collection of bacterial isolates from prostatectomy tissue^36^, we identified and cultured a strain of *P. lymphophilum* (strain API-1) capable of steroid metabolism (**Fig. 3a, b; Extended Data Figure 8**). We speculate that this isolate could have derived from prostatic fluid, or it could have been present in the prostatic urethra. Strain API-1 was found to generate products from cortisol (363.2 *m/z*; RT 3.20 min) consistent with 11OHAD (303.2 *m/z;* RT 3.63 min) and 11β-hydroxytestosterone (11OHT, 305.2; RT 3.42 min) that co-migrate with authentic standards (**Fig. 3c**). While side-chain cleavage of cortisol may be considered the “gateway reaction” in the formation of androgens from glucocorticoids by the urinary tract microbiota, the conversion of the 17keto (AD, 11KAD) to 17β-hydroxy (T, 11KT) is required to increase the “androgenicity” of these metabolites due to substrate-specificity by nuclear AR. To our knowledge, this is the first report of steroid 17β-HSDH activity by bacteria isolated from the urinary tract, let alone from prostate tissue. We sequenced and assembled a 2.3 Mb genome of *P. lymphophilum* API-1 (**Fig. 3c; Supplementary Table 4**). The *desABE* operon was located, providing an enzymatic basis for the conversion of cortisol to 11OHAD. We then searched for candidate 17β-HSDH genes in *P. lymphophilum* API-1, and a single candidate (ILDKDCJM_00761) in the SDR family appeared most probable based on genomic context (monocistron), as we have observed for HSDHs previously^37^. This gene is annotated as a “3β-hydroxycholanate dehydrogenase”, suggesting that the enzyme metabolizes steroids. This gene was cloned, overexpressed and affinity purified from *E. coli* BL21(DE3)RIPL (**Fig. 3d**). The Strep-Tactin affinity purified 26.8 kDa recombinant enzyme (10 nM) was incubated with NADPH and 11OHAD and 11OHT formation was confirmed by LC/MS (**Fig. 3e, f**). Thus, we have discovered the first microbial 17β-HSDH in a bacterium isolated from prostate tissue. This 17β-HSDH gene is part of the steroid-17,20-demolase pathway in *P. lymphophilum* API-1, so we propose to name this gene *desG* (**Fig. 3g**).

**Figure 3.**
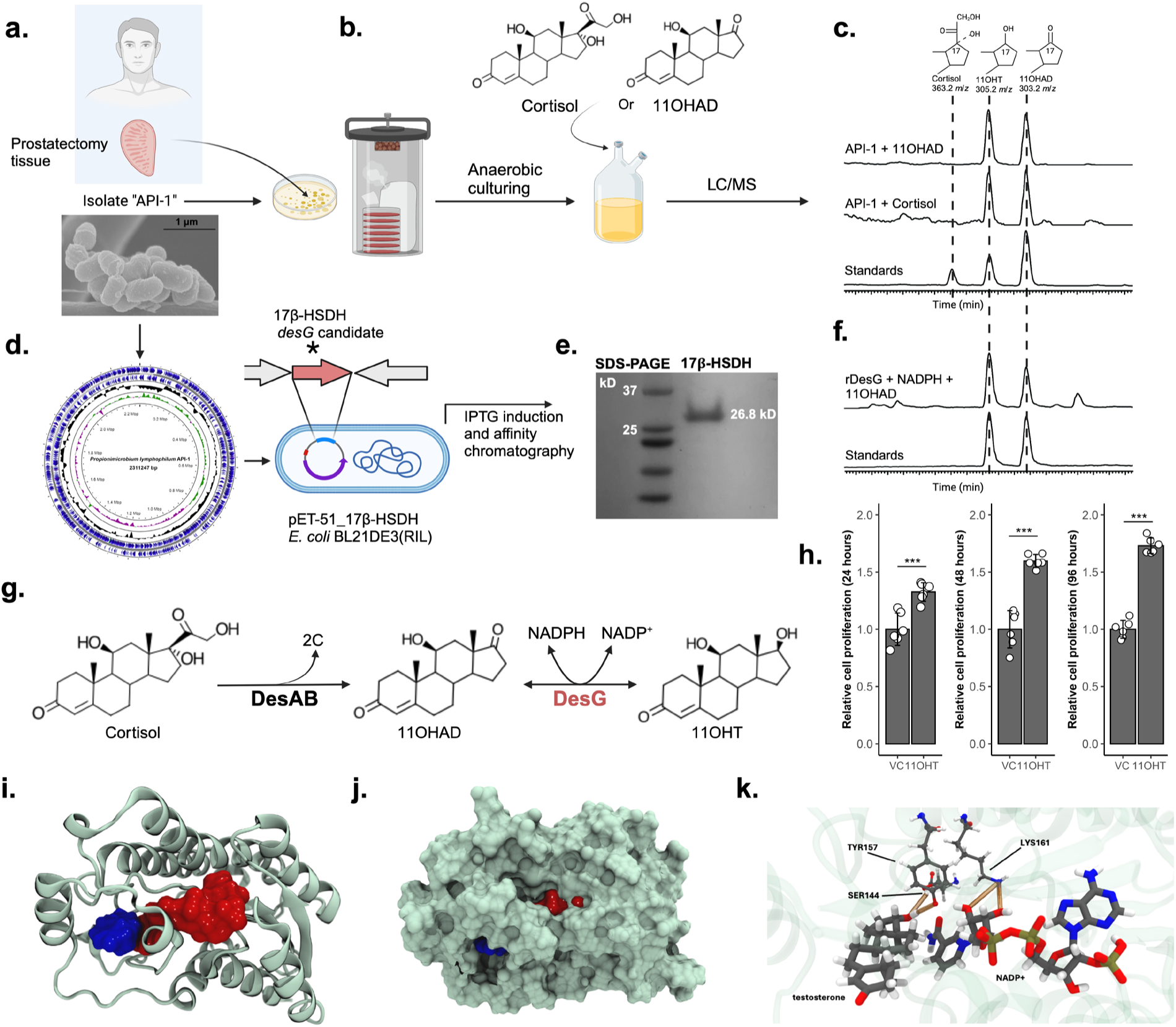
Androgen-producing bacterium isolated from human prostatectomy tissue expresses a multi-step pathway for conversion of glucocorticoids to derivatives of testosterone. **a-c.** A post-surgical prostatectomy specimen was placed in a sterile container following resection and the biopsy tissues were minced in sterile PBS, placed in a tube of thioglycolate broth, and cultured anaerobically at 37°C. A single colony capable of converting cortisol to 11OHAD and 11OHT was identified by LC/MS/MS. SEM imaging of the strain was obtained and the complete genome was sequenced. The isolate was identified as *Propionimicrobium lymphophilum* and was named “androgen-producing isolate 1” (API-1). **d.** A gene predicted to encode “3β-hydroxycholanate dehydrogenase” was identified in the genome sequence and selected as a potential 17β-HSDH candidate. **e.** We cloned ILDKDCJM_00716 into pET51b(+) and overexpressed and Streptactin-purified the recombinant protein for enzyme assay. **f.** LC/MS/MS analysis of steroid products after incubation of ILDKDCJM_00716, 200 µM NADPH and 50 µM 11OHAD. 11OHT formation was observed after 6 h incubation. We proposed the name *desG* for ILDKDCJM_00716. **g.** Pathway for conversion of cortisol to 11OHAD and 11OHT by *P. lymphophilum* API-1. **h.** The product of cortisol metabolism by strains API-1 and API-2, 11OHT, causes significant (*P < 0.001*) and prolonged (96 h) proliferation of LNCaP cells relative to vehicle control (VC; methanol 0.5% v/v). **i.** Ribbon diagram of AlphaFold 2 structural prediction of DesF. NADP^+^ (red) and testosterone (blue) are depicted as space-filling models and were aligned and placed with VMD and ligand-structure interactions were minimized using NAMD through its QuikMD interface. **j.** Space-filling model of both the DesG (gray) and ligands NADP^+^ (red) and testosterone (blue). **k.** Molecular dynamic trajectory analysis revealed strong interactions between ligands and catalytic triad. SER144 and TYR157 form stable hydrogen bonds with testosterone, while LYS161 was identified as key in NADP^+^ stabilization. Cell culture data are mean ± S.D. from six biological replicates. *P* values were calculated by unpaired *t*-test and Benjamini-Hochberg correction, * *P <* 0.05, ** *P <* 0.01, *** *P <* 0.001.

Using VMD, the structure of DesG was predicted by AlphaFold 2 and aligned with similar SDR PDB structures, as we had done with DesF, allowing us to minimize the ligand structure of NADP^+^ into the predicted binding pocket. The structure of testosterone in DesG was also fitted to the most probable binding site using VMD. MD simulation determined that in contrast to the open pocket observed with DesF, the results indicated that testosterone (blue) and NADP+ (red) occupy a pocket that closes over the catalytic region in DesG (**Fig. 3i, j**). In DesG, LYS161 is predicted to play a role in stabilizing NADP^+^, while SER144 and TYR157 formed stable interactions with the testosterone molecule (**Fig. 3k**). Simulations indicated that the predominantly hydrophobic clefts in both DesF and DesG are crucial for maintaining the stability of the complex, with steroid molecules fluctuating minimally in the cleft. Based on the MD simulations, we propose that the serine and tyrosine residues stabilize the steroid to initiate the enzymatic reactions, while the lysine residue helps hold NADP^+^ in place in both DesF and DesG.

We then performed RNA-Seq analysis in the presence of cortisol (n=5) or 11OHAD (n=5) vs. DMSO vehicle control (n=5) with *P. lymphophilum* strain API-1. We did not observe differential gene expression between vehicle control and cortisol treatment (50 µM) with respect to *desA* (ILDKDCJM_00614; 0.73 log_2_FC; 0.193 FDR), *desB* (ILDKDCJM_00613; 0.81 log_2_FC; 0.107 FDR), or *desG* (ILDKDCJM_00761; 0.28 log_2_FC; 0.61 FDR) (**Supplementary Table 5**). These results indicate that expression of steroid-17,20-desmolase genes in *P. lymphophilum* are not regulated by steroids, as is the case in the GI tract^6^, but rather the genes are constitutively expressed. This form of regulation may be important in the urinary tract due to the nanomolar levels of urinary steroids^18^ whose quantities are likely insufficient for sensitive inducible systems to evolve.

Protein phylogeny of the amino acid sequence of DesG revealed other taxa isolated in the urinary tract, but whose steroid metabolism remains unknown (**Extended Data Fig. 9**). Such sequences, which display >77% ID with the DesG from *P. lymphophilum*, include *Arcanobacterium urinimassiliense* (BQ7117_RS04815)*, Vaginimicrobium propionicum* (CZ356_RS01445), and *Sutterella wadsworthensis* (HMPREF1255_RS00895). Phylogeny and SSN analysis of DesAB from the gut bacterium *Csci*35704 led to identification of *A. urinimassiliense* as a *desAB* harboring urinary bacterium^3042^. To functionally sample from nearby neighbors of DesG, we cloned the synthesized genes predicted to encode 17β-HSDH in these species into pET51b(+) and expressed these recombinant proteins in *E. coli* BL21(DE3) (**Extended Data Fig. 10a-c**). After protein purification (**Extended Data Fig. 10c**) and incubation of 10 nM enzyme with 200 µM NADPH and 50 µM AD, the only protein that yielded T was rBQ7117_RS04815 from *A. urinimassiliense* (**Extended Data Fig. 10d)**. Together with our current phylogenetic results, this finding indicates that strains of *A. urinimassiliense* are also capable of generating 11OHAD and 11OHT from cortisol, along with strains of *P. lymphophilum* reported in this study. These data confirm a two-step pathway for converting glucocorticoids into 11OHT (**Fig. 3g).** To determine androgenicity of 11OHT, we measured time-dependent proliferation of LNCaP cells. Compared to VC, 10 nM 11OHT caused significant (*P <0.001*) and prolonged (96 h) growth of LNCaP cells indicating that this microbial pathway in the urinary tract generates androgens (**Fig. 3h**). Similar proliferation of VCaP cells was observed in the presence of 10 nM 11OHT (2 d, *P* = 1.9 × 10^-08^; 4 d, *P* = 1.3 × 10^-08^; 8 d, *P* < 2.0 × 10^-16^) relative to VC (**Extended** Data Fig. 6).

We then obtained a urine sample from the same male patient from whom we isolated *P. lymphophilum* API-1 from prostatectomy tissue to determine long-term colonization by strains similar to or evolved from *P. lymphophilum* API-1. This urine sample was collected approximately 17 years after prostatectomy (**Fig. 4a**). Upon isolating colonies from the urine sample, we screened for cortisol metabolism and obtained a bacterium capable of producing both 11OHAD and 11OHT (**Fig. 4b**). We sequenced the complete 2.1 Mb genome of this isolate and named it *P. lymphophilum* API-2 (**Supplementary Table 4**). Comparative genome analysis indicated an average nucleotide identity (ANI) of 98.9% to API-1 with both strains sharing 72.0% of their predicted protein coding genes (**Fig. 4c, d**). These results indicate that some individuals experience long-term colonization of androgen-producing urinary tract bacteria.

**Figure 4.**
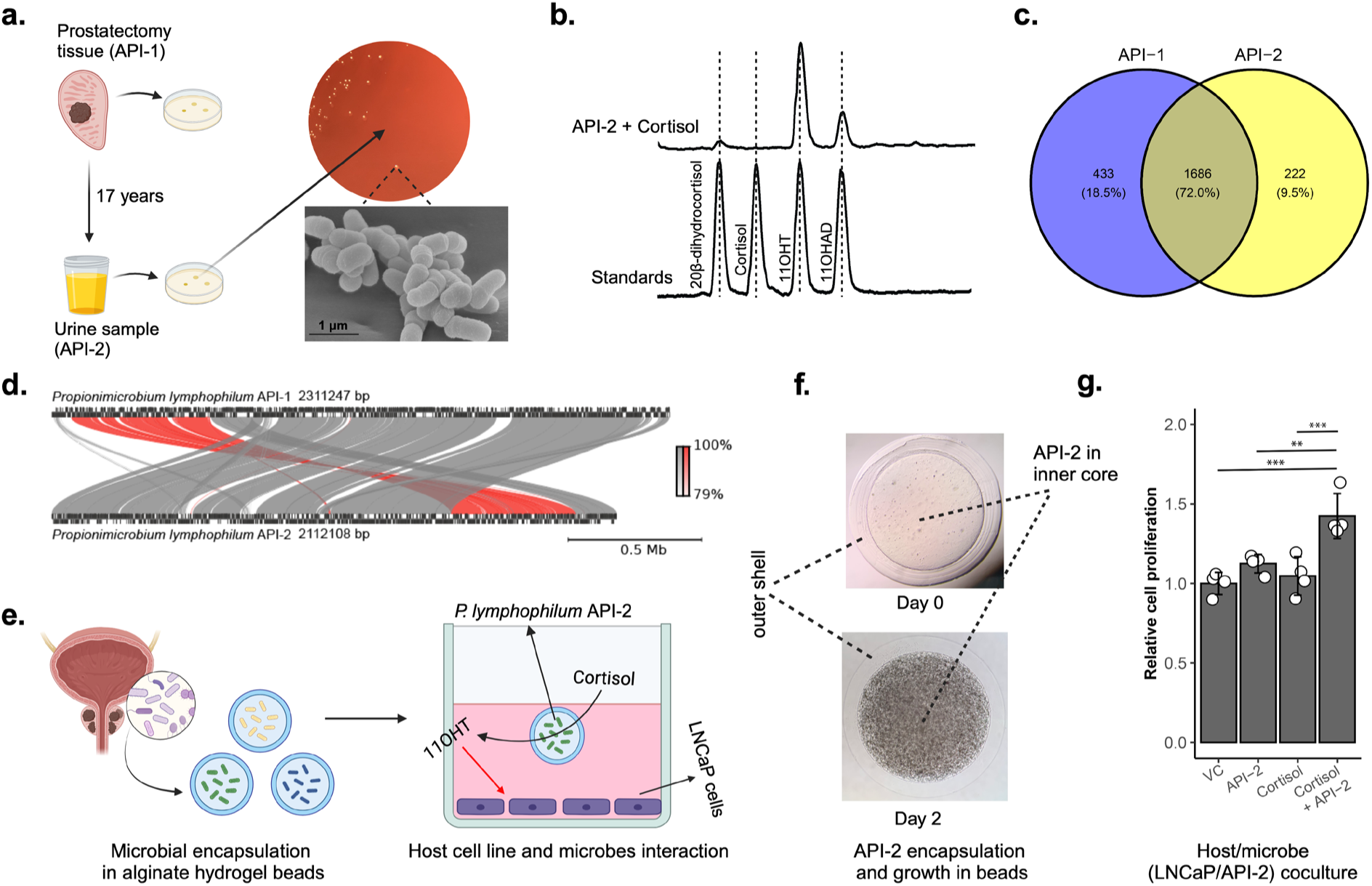
Urinary tract isolates of *Propionimicrobium lymphophilum* drives prostate cancer cell growth through androgen-production. **a.** Strain API-1 was isolated from prostatectomy tissue from a male with mCRCP, and strain API-2 was isolated ∼17 years later from a urine sample collected from the same patient. **b.** LC/MS analysis confirmed the conversion of cortisol to 11OHAD and 11OHT in encapsulated beads (see **e, f**). **c,d.** Synteny and comparison of gene content between the two strains indicates a high degree of similarity**. e.** Schematic representation of urinary tract isolates encapsulated in microgels which are co-cultured in culture medium with LNCaP cells in the presence of cortisol. **f.** Micrographs of calcium alginate microgels at Day 0 and Day 2 display dense growth of API-2 in DMEM medium under aerobic conditions. LC/MS analysis of cortisol metabolites extracted from DMEM in which API-2 was grown in microgels. **g.** LNCaP proliferation in the presence/absence of API-2 and/or 10 nM cortisol (n=4). Data are mean ± S.D. from four independent experiments *P* values were calculated by unpaired *t*-test and Benjamini-Hochberg correction, * *P <* 0.05, ** *P <* 0.01, *** *P <* 0.001.

### Androgen producing urinary microbes promote prostate cancer cell proliferation

We recently developed and reported a microencapsulation technique using calcium alginate beads to co-culture bacteria and host cells allowing metabolic interaction without direct contact (**Fig. 4e**)^38,39^. This bioengineered platform was utilized to determine the effect of cortisol metabolism on LNCaP cell proliferation. We first established the growth of *P. lymphophilum* API-2 in this platform in both anaerobic bacterial growth medium (PYG broth) (**Fig 4f**). LNCaP cell proliferation was determined in the co-culture platform in the presence of the bead encapsulated strain API-2 +/- 10 nM cortisol. Cells (50,000 cells/well) were seeded in a 24-well plate in RPMI 1640 media containing 10% charcoal stripped FBS. Cell proliferation assay was carried out after 96 h. Significant proliferation was observed only in the presence of strain API-2 and cortisol (**Fig. 4g**). These data confirm that the metabolism of cortisol by prostatectomy tissue/urinary tract isolates promote prostate cancer cell growth by generating androgen metabolites capable of activating AR-signaling.

### Culturomics of androgen-forming taxa in male human urine

To identify additional androgen-forming urinary microbial taxa, we collected clean catch urine from 25 patients during a pre-biopsy visit to Carle Hospital Oncology, and clean catch urine from 14 age-matched healthy controls (**Supplementary Table 6**). We first screened urine samples for conversion of cortisol to 11OHAD (*desAB* activity) or 11OHT (*desAB* and *desG* activity) by diluting freshly collected urine in PYG broth. LC/MS analysis identified 11OHAD and/or 11OHT production in urine from 8 of 25 pre-biopsy samples, 4 of which were subsequently diagnosed with prostate cancer, and 2 of 14 healthy control samples (**Fig. 5a**; **Supplementary Table 6; Extended Data Fig. 8**). Of the samples collected, 10 urine samples tested positive for 11OHAD and/or 11OHT production were plated (100 µL) on anaerobic Blood agar, Columbia agar and Schaedler agar. Colonies were cultivated in PYG broth in 96-well plates in the presence of 11DC in order to screen for *desAB* function, or 11OHAD to screen for *desG* function (**Fig. 5a**).

**Figure 5.**
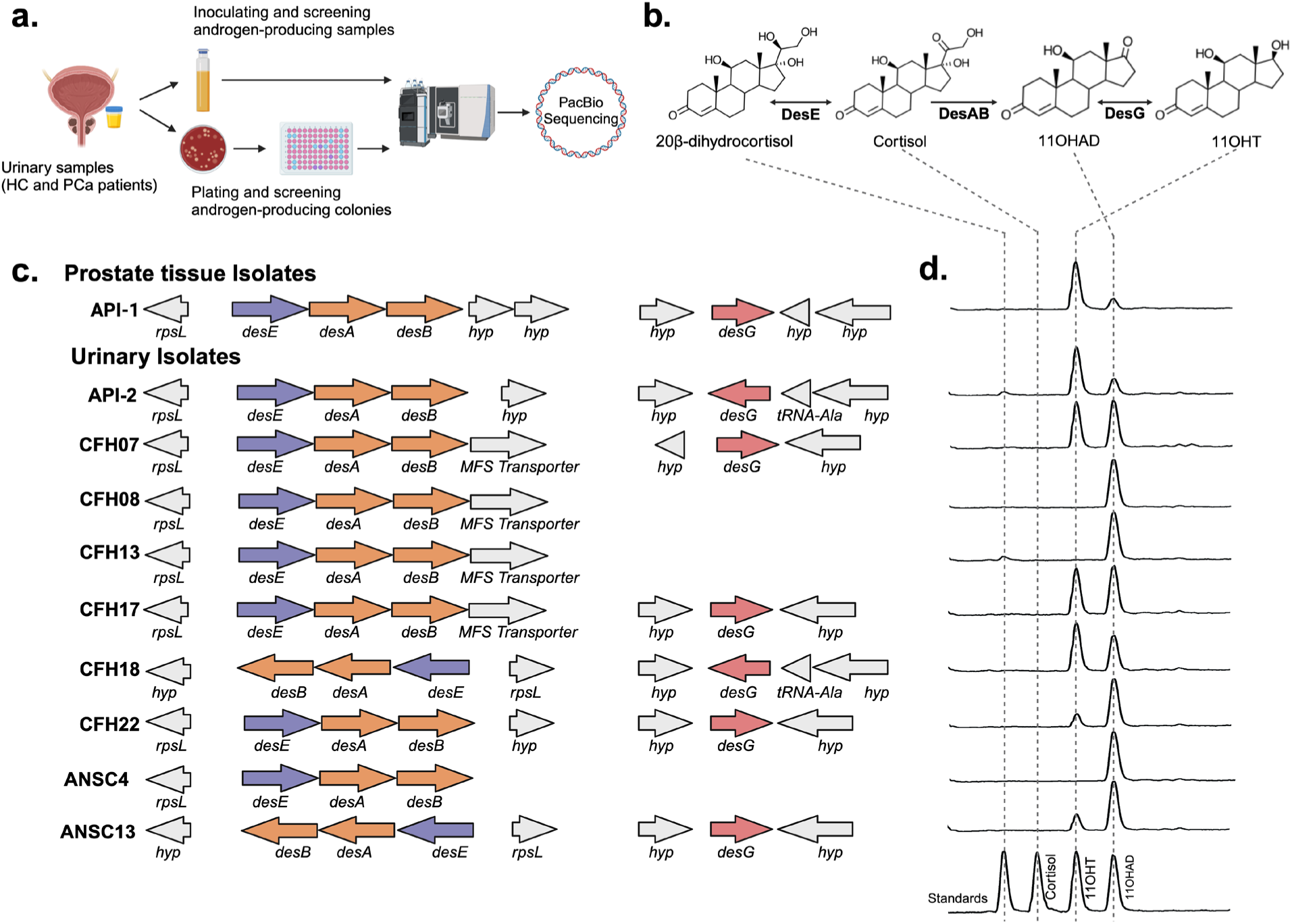
Isolation and characterization of androgen-producing bacteria from human male urine samples. **a.** Schematic of isolation and screening approach from human male urine **b.** Biochemical pathway and DesABEG enzyme functions proposed for urinary tract isolates. **c-d.** Organization of *desABE* and *desG* genes corresponds with the formation of 11OHAD and 11OHT in pure cultures incubated with 50 µM cortisol.

We obtained 9 *P. lymphophilum* isolates from urinary samples and sequenced their genomes (**Extended Data Figs. 8, 11)**. We identified the *desABE* genes encoding bacterial desmolase (*desAB*) and steroid 20β-HSDH (*desE*)^37,40^ in all strains of *P. lymphophilum* (**Fig. 5c**). Strikingly, 6 of 9 strains of *P. lymphophilum* also had 17β-HSDH activity involved in conversion of 11OHAD to 11OHT (**Fig. 5c**). We confirmed that the *desG* gene was found in the genomes of *P. lymphophilum* strains with both desmolase and 17β-HSDH activities, but not in the genomes of desmolase positive strains without 17β-HSDH activity (**Fig. 5c**). The *desABE* and *desG* genes and their metabolic activities in strain API-2 are consistent with that in strain API-1 (**Fig. 3g**; **Fig. 5c, d**).

### Abiraterone does not inhibit androgen production by bacterial desmolase

Host steroid-17,20-desmolase is encoded by *CYP17A1* (the drug target of AA) and functions as an NADPH and O_2_-dependent P450 monooxygenase that facilitates adrenal corticosteroid and androgen biosynthesis through the side-chain cleavage and 17ɑ-hydroxylation of pregnenolone^19^. CYP17A1 is inhibited by AA to treat prostate cancer through the inhibition of adrenal androgen synthesis. By contrast, bacterial steroid-17,20-desmolase (DesAB) functions under anaerobic conditions, through a predicted vitamin B1-dependent manner^7^ and may continue to function in the presence of AA, potentially contributing androgens from cortisone or P derivatives that may reduce the efficacy of AA/P therapy. We therefore tested AA (IC_50_ lyase activity = 15 nM) for inhibition of the bacterial desmolase (DesAB) by pre-treating early log-phase cultures of *P. lymphophilum* API-1 in PYG broth containing cortisol or prednisone. We determined, in two independent labs, that therapeutic and physiologically relevant concentrations of AA do not inhibit bacterial desmolase from generating androgens from glucocorticoids (**Extended data Fig. 12a, b**). Moreover, the active form of A was not able to inhibit DesAB activity (**Extended data Fig. 12c**).

## Discussion

This study significantly advances our understanding of the genetic potential of host-associated microbiota to produce androgens from glucocorticoids and androstenedione. We identified new genes (*desF, desG*) that expand the steroid-17,20-desmolase pathway that previously included only the side-chain cleavage enzyme (DesAB) and the side-chain oxidoreductases 20ɑ-HSDH (DesC) and 20β-HSDH (DesE), which we reported previously^6,7,37,40^. Specifically, our results reveal for the first time that (1) a novel bacterial pathway for conversion of AD (or cortisol derivatives) to epiT is encoded in the gut microbiome; (2) the end product, epiT, activates AR-dependent growth of LNCaP cells on par with T, indicating that epiT is a currently unrecognized AR agonist; (3) AA inhibits host production of adrenal androgens (AD, DHEA, 11OHAD) but not bacterial desmolase (*desAB* genes); (4) the *desF* gene is enriched in the fecal microbiota of individuals with advanced prostate cancer with disease progression on AA/P; and (5) enzymes in the epiT pathway may represent therapeutic targets for treatment of prostate cancer in some individuals in the same manner as the host steroidogenic enzymes are for drug targets. The discovery of the *desABC* and *desF* genes will allow direct quantification of microbial steroidogenic pathway genes that will complement fecal 16S rRNA profiling that can at most identify the abundance of *“C. scindens*”, strains which vary considerably in their capacity to metabolize steroids.

Moreover, we demonstrate for the first time that urinary tract bacteria, including a prostate tissue isolate, encode both *desAB* and the newly discovered *desG* gene that convert glucocorticoids (including prednisone) to T derivatives that promote prostate cancer cell proliferation. Urine is the main route of glucocorticoid excretion in humans, and glucocorticoids are measured in urine on the order of hundreds of nanomolar^19^. Based on this, we predict that it is possible that bacterial androgen production occurs locally in the prostatic urethra. How or whether this androgen production can influence the prostate is beyond the scope of the current study but suggests the need for future research. Intriguingly, studies have shown that *P. lymphophilum* abundance in urine is associated with prostate cancer^41,42^. We speculate that long-term colonization of the urinary tract by androgen-producing bacteria may be an underrecognized promoter of the development and/or progression of prostate cancer in some individuals (**Extended Data Fig. 13)**. Further clinical and mechanistic microbiome studies examining the role of androgen producing bacteria in the urinary tract in primary prostate cancer is warranted. The role of androgen producing bacteria in other human diseases driven by androgens in males such as benign prostate hyperplasia and in females such as breast cancer^43^ and polycystic ovary syndrome (PCOS)^44^ should also be examined.

## Materials and Methods

### Bacteria, cell cultures, and chemicals

#### Bacteria

*Clostridium scindens* ATCC 35704 (*Csci*35704) and *Clostridium scindens* VPI 12708 (*Csci*12708) were derived from in house 30% glycerol stock cultures and cultivated in anaerobic Trypticase Soy Broth (TSB) at 37°C. Individual colonies were picked from anaerobic TSB plates, DNA extracted, and identity confirmed by 16S rRNA gene sequence and confirmatory PCR targeting *baiJ* (*Csci*12708) or *desA* (*Csci*35704).

#### Cell culture

LNCaP and VCaP cells were obtained from ATCC and cultured in RPMI 1640 medium (Corning 10-040) and DMEM (ATCC 30-2002), respectively, and supplemented with 10% Fetal Bovine Serum (FBS) from GIBCO. Additionally, RPMI was supplemented with 10 mM HEPES buffer (Corning), 1 mM sodium pyruvate (Corning) and 4.5 g/L D-glucose (Sigma). For the androgen treatment experiments, cells were starved of androgens by cultivation in charcoal stripped FBS (GIBCO) medium. Cell passage number was kept below 25 for all experiments. Both cell lines were tested for mycoplasma contamination and authenticated using STR analysis in TEP CCIL.

#### Steroids

Steroid (commercial sources) included: cortisol (Sigma); 11β-hydroxy-androstenedione (11OHAD, Steraloids, Newport, RI, USA);11β-hydroxy-testosterone (11OHT, Steraloids); Epi-testosterone (epiT, Steraloids); 11-deoxycortisol (11DC, Sigma); Androstenedione (AD; Sigma); Testosterone (T, Sigma); 1,4-androstadiene-3,11,17-trione (AT, Sigma); 11-deoxycortisol-D5 (2,2,4,6,6-D5) (11DC-D5, Sigma); Androstene-3,17-dione-2,3,4-13C3 solution (AD-13C3, Sigma); Testosterone-D3 (16, 16, 17-d3) solution (T-D3, Sigma); 17-epi-testosterone-D3 (epiT-D3, Santa Cruz); Enzalutamide (Selleck); Abiraterone (A, MedChemExpress); Abiraterone acetate (AA, Sigma). T and AD were purchased in solution form (Sigma), evaporated with nitrogen, and redissolved in DMSO at the required concentration. EpiT, cortisol, 11OHAD, 11OHT, prednisone and AT were dissolved in DMSO or methanol. Enzalutamide was dissolved in DMSO.

### Bacterial media preparation

Brain Heart Infusion (BHI, BBL) broth was purchased and prepared based on the instructions. The Trypticase Soy Broth (TSB, BBL) was prepared as instructed with the addition of 5 g yeast extract, 1 g L-cysteine, 1 mg resazurin and 40 mL salt solution (1 L; 0.25 g CaCl_2_•2H_2_O, 0.5 g MgSO_4_•7H_2_O, 1 g K_2_HPO_4_, 1 g KH_2_PO_4_, 10 g NaHCO_3_, 2 g NaCl). Peptone Yeast Glucose (PYG) broth (modified) was prepared according to the DSMZ protocol. Briefly, each liter contains 5 g trypticase peptone, 5 g peptone, 10 g yeast extract, 5 g beef extract, 5 g glucose, 2 K_2_HPO_4_, 1 mL tween 80, 40 mL salt solution (see above), 1 mg resazurin, 0.2 mL vitamin K1 solution (5mg/ml), 10 mL hemin solution (50 mg/100 mL), 1 g L-cysteine. Blood agar base (Sigma) was purchased and prepared according to the instructions with 6% (v/v) defibrinated sheep blood (Thermo Scientific) added. Schaedler agar (Sigma) was purchased and prepared as instructed. Columbia broth (BBL) was purchased and prepared as instructed with 15 g/L agar added to solidify the medium. The broth was made anaerobic by storage in Hungate tubes with 100% N_2_ in the headspace. Plates were made anaerobic by storage in an atmosphere of 85% N_2_:10% CO_2_:5% H_2_.

### Whole-cell steroid conversion assay

*Csci*35704 and *Csci*12708 were precultured in TSB. Afterwards, each fresh culture of these two strains was inoculated (0.5 mL) into fresh TSB with 50 µM 11DC. For sampling, 1-mL samples were collected at 0 and 24 h. The collected samples were clarified by centrifugation (13,300 × *g*, 10 min; Thermo Scientific) and the supernatant fluid was used for subsequent analysis.

*P. lymphophilium* strains isolated from prostatectomy tissue or urine samples were cultured in anaerobic PYG broth. Log-phase cultures were transferred to fresh PYG broth containing 50 µM cortisol and incubated for 48 h. These samples were extracted and analyzed by LC/MS for 11-oxyandrogen formation as discussed below.

### Steroid extraction

Two parts ethyl acetate and 1 part bacterial culture supernatant were thoroughly mixed by vortexing for 1 min. Next, the ethyl acetate layer was carefully collected and transferred to new tubes. The extraction process was repeated, and the collected top layers were evaporated with nitrogen gas and dissolved in 200 µL LC-MS grade methanol. For samples where substrates and end products were to be quantified, an internal standard mixture of 11DC-D5, AD-13C3, and epiT-D3 were added before extraction. The concentrations of 11DC, AD and epiT were normalized based on the under-curve area of the internal standards accordingly.

### Liquid chromatography-mass spectrometry (LC-MS)

Samples were sent to the Mass Spectrometry Lab (University of Illinois at Urbana-Champaign, Urbana, Illinois, USA) for metabolite analysis using liquid chromatography-mass spectrometry (LC-MS). LC-MS for all samples was done on a Waters Aquity UPLC coupled with a Waters Synapt G2-Si ESI MS (Waters Corp., Milford, MA, USA). Chromatography was performed using a Waters Acquity UPLC BEH C18 column (1.7 μm particle size, 2.1 mm x 50 mm) at a column temperature of 40°C and an injection volume of 0.5 µL. For gradient elution, 2 mobile phases were used: mobile phase A contained 95% water, 5% acetonitrile, and 0.1% formic acid; mobile phase B contained 95% acetonitrile, 5% water, and 0.1% formic acid. Initially, mobile phase A was 100% for 0.5 min. Over the next 5.5 min, mobile phase B linearly increased, reaching 70% at 6 min. Then, mobile phase B increased to 100% in 1 min and maintained for 1 min. Afterwards, a steep reversal to the initial conditions was done within 0.1 min, and the running condition was maintained until the end at 10 min. The flow rate was 0.5 mL/min. The LC eluents were introduced into the mass spectrometer equipped with electrospray ionization (ESI) with a positive ion mode for steroid analysis. The following optimized conditions were used: capillary voltage of 3 kV, desolvation temperature of 500 °C, cone voltage of 25 V, collision energy of 4 eV, collision gas helium, source temperature of 120°C, cone gas flow of 10 L/h, and desolvation gas flow of 800 L/h. The mass range was 50–2000 Da. Mass Lynx v4.1 (Waters) was used for chromatographs and mass spectrometry data analysis.

### Nuclear magnetic resonance (NMR)

AD metabolism was performed in *Csci*12708 cultures grown at 37°C in anaerobic BHI medium in the presence of 25 μM of AD. Following overnight incubation, growth was quenched by adding 2 x volume ethyl acetate. After vortexing for 1 min, the organic layer was collected and evaporated to dryness under N2 gas. Dried extracts were resuspended in 500 μL methanol. One hundred microliters were injected and run on high-performance liquid chromatography (HPLC, Agilent) using a C18 reverse phase column (Agilent Eclipse XDB-C18), with a 50:50 methanol:water mobile phase at a flow rate of 1 mL/min. The absorbance of steroid metabolites was monitored at 240 nm by UV-Vis detector spectroscopy. The *C. scindens* VPI 12708 androstenedione metabolite was fractionally collected and sent for NMR analysis. ^1^H and ^13^C-NMR spectroscopic data were obtained on a JNM-ECA800 (JEOL, Ltd., Tokyo, Japan) instrument operated at 800 and 200MHz, respectively, with CDCl_3_ as the NMR solvent. Chemical shifts were expressed in d (ppm), and coupling constants *J*_H,H_ are given in Hz. ^1^H–^1^H nuclear overhauser effect spectroscopy (NOESY), ^1^H-^1^H correlation spectroscopy (COSY), ^1^H-^13^C heteronuclear single-quantum correlation spectroscopy (HSQC), and ^1^H-^13^C heteronuclear multiple-bond correlation spectroscopy (HMBC) spectra were obtained using gradient-selected pulse sequences. The ^13^C distortionless enhancement by polarization transfer (135°, 90°, and 45°) spectra were measured between CH_3_, CH_2_, CH, and coherence based on their proton environments.

### Computational search of reductases unique to *Csci*12708

A total of 261 *Csci*12708 protein sequences (CP113781.1) annotated as dehydrogenases, reductases and NADH dependent proteins, were subject to screening with the NCBI/CDSEARCH (https://www.ncbi.nlm.nih.gov/Structure/cdd/wrpsb.cgi), looking for short chain dehydrogenases/reductases, medium chain reductases, and aldo/keto reductases family candidates. NCBI/BLAST was used to filter unique protein sequences by removing any hit found above ∼95% identity. The remaining 38 candidate’s domain architecture was visualized with EMLB-EBI/InterPro (https://www.ebi.ac.uk/interpro/). Theorical molecular weight was estimated with Expasy algorithms (https://www.expasy.org/resources/compute-pi-mw). Similarity within sequences was tested with EMLB-EBI MUSCLE/Clustal2.1 (https://www.ebi.ac.uk/jdispatcher/msa/muscle).

### RNA-Seq analysis

*Csci*12708 was incubated in BHI broth in the presence/absence of 50 μM of 11OHAD at 37°C for 24 h. Cultures (10 mL) were pelleted by centrifugation (4,000 x *g*). RNA was extracted as previously described^6^. Samples were sent to Roy J. Carver Biotechnology Center, DNA Services Laboratory (University of Illinois at Urbana-Champaign, Urbana, Illinois, United States) for library construction and sequencing. Total RNAs were run on a Fragment Analyzer (Agilent) to evaluate RNA integrity. The total RNAs were converted into individually barcoded polyadenylated RNAseq libraries with the Kapa HyperPrep mRNA kit (Roche, CA), with prior removal of rRNAs with the FastSelect Bacteria kit (Qiagen, CA). Libraries were barcoded with Unique Dual Indexes (UDI’s) which have been developed to prevent index switching. The adaptor-ligated double-stranded cDNAs were amplified by with the Kapa HiFi polymerase (Roche). The final libraries were quantitated with Qubit (ThermoFisher) and the average cDNA fragment sizes were determined on a Fragment Analyzer. The libraries were diluted to 10 nM and further quantitated by qPCR on a CFX Connect Real-Time qPCR system (Biorad) for accurate pooling of barcoded libraries and maximization of number of clusters in the flowcell. The barcoded RNAseq libraries were loaded on one SP lane on a NovaSeq 6000 for cluster formation and sequencing. The libraries were sequenced as single-reads 100nt in length. The fastq read files were generated and demultiplexed with the bcl2fastq v2.20 Conversion Software (Illumina, San Diego, CA).

Read quality was evaluated using FastQC v0.11.8^45^. SeqKit v2.0.0^46^ was used to calculate the read number, sum of the read length, minimum read length, average read length and maximum read length for each sequencing file. Trimmomatic v0.39 was used to remove the adaptors and low-quality reads^47^. SortmeRNA v4.3.6 was used to filter out ribosomal RNAs^48^. Salmon v0.14.1 was used to do gene quantification^49^ using *C. scindens* VPI 12708 genome as a reference^28^. Gene abundance was filtered and normalized using the edgeR package^50^. Differential gene expression analysis was performed using the limma package^51^.

### Genomic DNA isolation

Genomic DNA of *C. scindens* ATCC 35704, *C. scindens* VPI 12708 and *P. lymphophilum* API-1 were extracted using the QIAamp PowerFecal Pro DNA kit (Qiagen) according to the manufacturer’s instructions. The extracted DNA was used to amplify the target genes for the heterologous expression of the potential candidates.

### Heterologous expression and purification of potential 17α-HSDH and 17β-HSDH proteins

The target inserts were amplified using the primers (**Supplementary Table 7**) synthesized by Integrated DNA Technologies (IDT, Coralville, IA, USA) and Phusion High Fidelity Polymerase (Stratagene, La Jolla, CA, USA). Inserts and the pET-51b(+) plasmid (Novagen, San Diego, CA, USA) were double digested using the appropriate restriction enzymes (**Supplementary Table 7**) (NEB, Ipswich, MA, USA). The digested inserts and plasmids were ligated using the T4 DNA ligase. Recombinant plasmids were transformed into *E. coli* DH5α cells via heat shock at 42°C for 30 seconds. *E.coli* DH5α were plated on lysogeny broth (LB) agar plates supplemented with 100 µg/mL ampicillin. Single colonies were picked up and transformed into 10 mL LB broth with 100 µg/mL ampicillin and grown for 10 h prior to plasmid extraction from cells. The plasmids were extracted using the QIAprep Spin Miniprep kit (Qiagen, Valencia, CA, USA). The extracted plasmids were transformed into *E. coli* BL-21 CodonPlus (DE3) RIPL chemically competent cells by heat shock method as mentioned above and cultured overnight at 37°C on LB agar plates supplemented with 100 µg/mL ampicillin. Selected colonies were precultured in LB broth with 100 µg/mL ampicillin for 6 h at 37°C, and subsequently added to fresh LB medium (1 L), supplemented with 100 µg/mL ampicillin. When the OD_600nm_ reached 0.4, the incubation temperature was decreased to 25°C and IPTG was added to each culture at a final concentration of 0.1 mM to induce the protein production duing the 16-h incubation.

Subsequently, cells were pelleted and resuspended in 30 mL of binding buffer (10 mM Tris, 400 mM NaCl, 10 mM 2-mercaptoethanol, pH 8.0). To lyze cells, 750 μl of lysozyme (1 mg/mL) and 10 μl of benzonase nuclease (Sigma) were added and incubated on ice for 40 min. Afterwards, cells were physically lysed by passing through a French pressure cell press twice. Cell lysate was separated by centrifugation (13,300 × *g*) at 4°C for 30 min. The recombinant proteins in the soluble fraction were purified using Strep-Tactin resins (IBA lifesciences) according to the manufacturer’s instructions. The purified proteins were assessed by sodium dodecyl sulfate-polyacrylamide gel electrophoresis (SDS-PAGE). Protein concentrations were measured by Nanodrop 2000c spectrophotometer based on their extinction coefficients and molecular weights. Purified proteins were sent to Roy J. Carver Biotechnology Center, Proteomics Core (University of Illinois at Urbana-Champaign, Urbana, Illinois, United States) for protein confirmation and size determination.

### Enzyme assays

Purified recombinant 17α-HSDH and 17β-HSDH activities were determined by mixing 10 nM enzyme, 50 μM substrate, and 200 μM cofactor (NADPH/NADP^+^) in phosphate-buffered saline. Samples were collected before and after the reactions. Steroids were extracted as mentioned above. Extracted samples were sent to the Mass Spectrometry Lab (University of Illinois at Urbana-Champaign, Urbana, Illinois, United States) for metabolite analysis using LC-MS.

### Proteomics

Samples of affinity purified DesF were digested with Trypsin (Thermo) using a CEM Discover Microwave Digestor (Matthews, NC) at 55⁰C for 30 min. The digested peptides were extracted, lyophilized, and cleaned up using stage-tips^52^. LC/MS was performed using a Thermo Fusion Orbitrap mass spectrometer in conjunction with a Thermo RSLC 3000 nano-UPLC. The column used was a Thermo PepMap C-18 (0.75 mm x 25 cm) operating at 300 nL/min and 40⁰C. The gradient was from 1% to 35% acetonitrile + 0.1% formic acid over 45 min time interval. The mass spectrometer was operating in positive mode using data dependent acquisition method and collision induced dissociation for fragmentation at 35% energy. The raw data were analyzed using Mascot (Matrix Science, London, UK) and searches were made using a database consisting of all the *desF* and *desG* sequences. FDR (False Data Discovery Rate) using a decoy reversed sequence database was at 1%.

### Sample collection and prostate cancer study cohort

All specimens were studied under a Johns Hopkins Medicine Institutional Review Board (IRB) approved protocol with written informed consent. Study participants (n=44) were instructed to collect a full stool sample followed by self-collection of rectal swabs (FLOQSwabs, Copan). The majority of the fecal samples used in this study were derived from rectal swabs. Seven of the fecal samples were swabs of stool. The samples were stored at −80°C until time of DNA isolation.

All of the study participants had advanced prostate cancer and were undergoing treatment with AA/P. The majority of participants had castration resistant disease (**Supplementary Table 3**). Samples were categorized as “stable” (n=28) if the donor had circulating PSA levels at the time of sample collection that were decreasing or had not changed from the prior PSA measurement. Samples were categorized as “progressing” if the donor had circulating PSA levels that had increased at least 0.2 ng/mL from the nadir, and that continued to rise (n=27) or that was re-detectable after a prolonged period of being below the limit of detection (n=1). Twelve participants in the study had matched fecal samples collected while stable on AA/P and then while progressing on AA/P.

### Fecal DNA isolation and *desF* quantitative PCR (qPCR)

DNA was isolated from fecal samples as previously described^53^. Samples were diluted to 10 ng/µL in DNA-free water. For each 20 µL reaction, the following reagents were combined: 10 µL of iQ SYBR Green Mix (Cat No. 1708882, Bio Rad Laboratories), 2 µL of 10 µM Forward/Reverse Primer set, 6 µL DNA-free water, 1 µL of 2 ug/µL BSA (Cat No. B14, ThermoFisher Scientific), and 1 µL of 10 ng/µL DNA. Real-time PCR (qPCR) conditions and primers are outlined in **Supplementary Table 8**. Total copies of *desF* were estimated using standard curves with genomic DNA extracted from *Csci*12708. The qPCR efficiency of all qPCR assays was determined to be between 86-114%.

### *Csci*12708 standard curve calculations

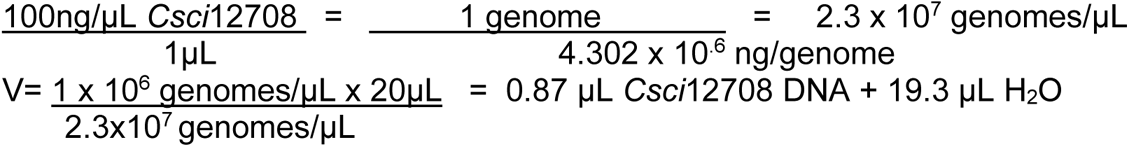

A serial dilution was made from this stock solution as standard 1 down to standard 6 (1×10^6^ to 1×10^2^). All qPCR data were verified by running the PCR products on a 1.5% agarose DNA gel and confirmation of a band at the correct size.

### Optimization of the *desF* primer set

To optimize the annealing temperature for the *desF* primer set, we ran a gradient annealing temperature plate with a positive control of genomic DNA from *Csci*12708, and negative controls of genomic DNA from *Csci*35704, the ATCC 20 strain mix (Cat No. MSA-1002, ATCC), human prostate DNA, and a no template control. Three gradient annealing temperature qPCR’s using the above samples were run for a range of 52-54.9°C, 56.1-60°C and 61-64°C. An annealing temperature of 65°C was chosen based on the gradient PCRs and the positive and negative controls. The optimized cycling conditions were run with the positive control, negative controls, and a *Csci*12708 standard curve to confirm correct melt curve data and adequate qPCR efficiency. Lastly, the protocol was tested with 10 ng of *Csci*12708 combined with 10 ng of human prostate DNA, to ensure that human background DNA would not impact the specificity of the protocol towards bacterial genomic DNA in the clinical samples. A qPCR utilizing the *Csci*12708 standard curve both in the presence and absence of human prostate DNA was run. Both standard curves resulted in the same data, indicating that human background DNA would not impact the results.

### Isolation of *P. lymphophilum* API-1 from prostatectomy tissue

The post-surgical prostatectomy specimen was placed in a sterile container following resection and transported to the grossing room. Here, a sterile field was assembled under a vertical laminar flow module (Envirco Corporation) for collection of tissue cores. A Biopty gun and sterile, single-use Biopty needles (18 gauge x 16 cm, C.R. Bard) were used to obtain two cores from both the right and left lobes of the prostate. Biopsy needles were positioned from apex to base, sampling the posterior (peripheral) aspect of the prostate. The biopsy tissues were minced in sterile PBS, placed in a tube of thioglycolate broth, and cultured anaerobically at 37°C. Stock cultures were stored in 33% glycerol at −80°C.

### Patient recruitment and urine sample collection (CCIL study)

To identify androgen-forming urinary microbial taxa, we consented and recruited 25 patients and 14 healthy individuals under the IRB #22383 (UIUC), Carle Hospital #18CCC1757. **Inclusion criteria**: No history of PCa, aged >50 yrs and <90. A BMI <35. Not having active diabetes. **Exclusion criteria**: Patients currently being treated for sexually transmitted infection, or urinary tract infection. Any patient taking antibiotics within the last month. Currently being treated for benign prostatic hyperplasia (BPH) i.e. with drugs such as alfuzosin (Uroxatral), doxazosin (Cardura), tamsulosin (Flomax), and terazosin (Hytrin) or abiraterone + prednisone or similar drugs. Unable or unwilling to give informed consent. Participants were given a urine collection kit and detailed instructions on how to properly collect a “clean/sterile catch”. The urine collection kit contained an alcohol swab to clean the urethra and tip of the penis. Participants were directed to catch mid-stream urine in sterile vials. At least 20 mL of urine was collected from participants. All urine samples were labeled with a unique, non-identifying code and were not derived from, or related to the participant’s personal information. The urine samples were processed immediately for culturomics work.

### Urine culturomics

To test urine samples for androgen-producing bacteria, 100-200 µL urine from each patient was screened in PYG broth containing 11DC and 11OHAD as substrates. 11DC was used to confirm side-chain cleavage activity (desAB) followed by 17β-HSDH activities in the urine yielding AD and T whereas 11OHAD conversion to 11OHT can identify 17β-HSDH activity when desAB-encoding microbes are lacking in the sample. LC/MS was performed to analyze metabolism of the substrates in these cultures.

Urine (100 µL) from each individual was plated (duplicate plates for each agar) to Blood agar, Columbia agar and Schaedler agar respectively. Each plate from each agar was incubated aerobically or anaerobically at 37°C. After 4-5 days, single colonies were picked using sterilized toothpicks to a 96-well plate containing PYG broth supplemented with 50 µM 11DC and 11OHAD. After incubation in an anaerobic chamber for 5 d, each column (50 µL/well) of 96 well plate was pooled to a single 1.5 mL centrifuge tube and extracted for the steroid. Columns positive for the steroid conversion were further processed to identify the individual positive well and the microbe carrying the conversion activity.

### Bacterial colony morphology and scanning electron microscopy (SEM)

Androgen-producing bacterial isolates were cultured in PYG broth in an anaerobic chamber. During the log phase, these broth cultures were streaked on blood agar plates to determine colony morphology. For colony morphology, the blood agar plates were imaged under the Amscope microscope. Karnovsky’s fixative, containing 2% glutaraldehyde and 2.5% paraformaldehyde was used to fix the bacterial isolates. Cultures (500 µL) in log phase were mixed with fixative (500 µL), vortexed, and then centrifuged 4,000 x *g* for 5 min at 4°C. Supernatant was discarded, and the pellet was washed 3 times with fixative (500 µL) and then stored at 4°C. Samples were sent to Materials Research Laboratory Central Research Facilities (University of Illinois at Urbana-Champaign, Urbana, Illinois, United States) for SEM imaging.

### Cortisol metabolism by *desAB* encoding bacteria in the presence of abiraterone (A) and abiraterone acetate (AA)

Abiraterone binds irreversibly to CYP17A host enzyme and inhibit pregnenolone and progesterone conversion to AD and T. *P. lymphophilum* API-1 was cultured in an anaerobic chamber for 3 days. These cultures were then treated with A (50 µM) or VC (DMSO) to find the inhibitory effect of A on *desAB* enzyme activity. After 24 h, cortisol (50 µM) was added to cultures and incubated for 72 h. At the end, 200 µL cultures were extracted with ethyl acetate (as described above); extracts were dissolved in LC-MS grade methanol and subjected to LC/MS. In a separate study, API-1 was grown in RCB media under anaerobic conditions. 500 µL of API-1 + 50 µM of steroid (cortisol or prednisone) + 1 uM of abiraterone acetate was added to a 7.5 mL RCB tube. This tube was grown anaerobically for 48 hours. The bacteria were then pelleted by centrifugation at 15,000 x g for 3 min. The supernatant was collected and used for LC/MS/MS that was performed at the Analytical Pharmacology Shared Resource at Johns Hopkins University.

### Cell proliferation assay

Androgen-starved LNCaP and VCaP cells were trypsinized and seeded in 96-well plates at a density of 10,000 cells/well and then treated with enzalutamide (2 µM) or VC (DMSO; final concentration 0.2%). After 24 h, these cells were treated with VC (0.5% methanol) or 10 nM T, epiT, AD, and 11OHT. There were 6 replicates for each treatment. 3-(4,5-dimethylthiazol-2-yl)-5-(3-carboxymethoxyphenyl)-2-(4-sulfophenyl)-2H-tetrazolium (MTS) reagent (15 µL) was added to each well at the end of the experiment and incubated for 90 min. Absorbance was measured at 490 nm in a Biotek Synergy HT plate reader.

To determine if epiAT derived from AT metabolism by *Csci*12708 impacted proliferation of LNCaP cells, *Csci*12708 was cultured in anaerobic TSB (10 mL) containing 50 µM AT or VC (DMSO). For sampling, 1 mL cultures were collected at 0, 24, 48 and 72 h. Metabolites of AT to epiAT by *Csci*12708 were analysed by LC-MS. The spent cultures (72 h) were filtered through 0.2 µm-syringe filters prior to adding to LNCaP cell cultures. The filtered (sterilized) spent culture fluids were divided into the following treatment groups: 1) VC (TSB-DMSO spent culture); 2) AT spiked (TSB-DMSO spent culture with 50 µM AT added); and 3) epiAT (TSB-AT spent culture; *Csci*12708 encodes the *desF* gene and converts AT to epiAT during growth). LNCaP cells were seeded to 96-well plates as mentioned above and exposed to the equivalent of 0.1, 1 and 10 nM of AT or epiAT from the above treatment group 2 and 3, respectively. After 4 d of incubation, cell proliferation assay was conducted to compare cell proliferation between different treatments.

### RNA extraction and gene expression qPCR from mammalian cells

LNCaP cells incubated in cRPMI medium for 24 h were seeded in 12-well plates at a density of 100,000 cells/well. Cells were treated with VC (DMSO) or 2 µM enzalutamide. After 24 h, cells were treated with T, epiT or VC and incubated for an additional 24, 48, and 96 h. Cells were trypsinized and pelleted in 1.5-mL microcentrifuge tubes by centrifugation (500 x *g*). Total RNA was extracted from cell pellets using GeneJET RNA Purification kit (ThermoScientific) where the lysis buffer was supplemented with 2% of 14.3 M β-mercaptoethanol. DNA contamination was removed from the extracted RNA using RapidOut DNA Removal Kit (thermoscientific). Total RNA was measured using Nanodrop and 100 nM high-quality RNA was converted to cDNA with High-Capacity cDNA Reverse Transcription Kit (Applied Biosystems, Thermofisher, Waltham, MA). Real-time PCR (StepOnePlus Real-Time PCR Systems; v 2.0 Applied Biosystems, Waltham, MA) was used to analyze differential gene expression. A total reaction volume of 20 µL in each well contained 0.5 µM forward and reverse primers (**Supplementary Table 8**), 6 ng cDNA and and PowerUp SYBR Green Master Mix (Applied Biosystems). Glyceraldehyde 3-phosphate dehydrogenase (GAPDH), a housekeeping gene was used as an exogenous reference to normalize transcription of PSA gene and the subsequent data was analyzed by the ΔΔCt method.

### Bacterial encapsulation and host/microbe coculturing

*P. lymphophilium* API-2 strain was cultured in PYG broth for 3 d in an anaerobic chamber. Encapsulation processes were followed by the reported method^54^. Briefly, bacterial cultures in the exponential phase were collected and then thoroughly mixed with 3 wt% (w/v) of sodium alginate (Sigma) solution to a final concentration of 2.5% alginate. Alginate core beads were fabricated by dropping alginate/bacteria mixture into a 100 mM CaCl_2_ solution while stirring for 10 min. The core beads encapsulating bacteria were transferred to a low concentration of alginate solution (less than 0.1 wt%) to fabricate a hydrogel-shell layer. The concentration of the alginate solution was increased very quickly by adding 3% of alginate solution to a final concentration of 0.5%. The reaction container was then vigorously shaken for 3 min to prevent core-bead aggregation. The process of shell fabrication was stopped by diluting the alginate solution when added an excessive amount of deionized (DI) water. The formed core–shell beads were transferred into a 0.01 M CaCl_2_ solution under mild stirring for stabilization then washed with DI water. Hydrogel beads were immediately transferred to the anaerobic chamber and incubated in PYG for 4 d.

For the coculture study, *P. lymphophilium* API-2 beads and control (empty beads) were washed with DI sterilized water and conditioned in cRPMI medium for 6 h. The bacterial and control beads were then transferred to the 24-well plate. LNCaP cells maintained in cRPMI were sub-cultured to 24-well plates (50,000 cells/well). LNCaP/API-2 coculture platform was treated with cortisol (final concentration 10 nM) or VC (5% methanol) and incubated. After 96 h, beads were carefully removed from the cells using a 10-µl inoculation loop. Cell proliferation assay was conducted by adding 50 µl MTS reagent to each well, incubating for 90 min, and reading the absorbance (490 nm) using a Biotek Synergy HT Plate Reader.

### High-molecular-weight DNA extraction

Urinary isolates were incubated at 37°C in anaerobic PYG broth (10 mL). During log growth, cells in cultures (10 mL) were harvested by centrifugation (4,000 rpm) at 4°C for 15 min). Cell pellets were washed 3 times with 1 mL TE buffer (10 mM tris, 1 mM EDTA, pH 7.6), and resuspended in TE buffer (450 μL) after the final washing. Bacterial lysis was carried out by the addition of lysozyme (2 mg/mL) with an incubation at 37°C for 1 h. Next, proteinase K (1 mg/ml) was added, and the samples were incubated at 56°C for 30 min. RNase A (4 µL, 100 mg/mL, Qiagen) was added to remove RNA and incubated at 25°C for 2 min. After that, sodium dodecyl sulfate (SDS) was added (1%, Sigma) and the samples were incubated at 60°C for 30 min. Equal volume phenol was added to denature proteins and then the samples were cleaned with 1 mL phenol/chloroform/isoamyl alcohol (25:24:1 vol/vol/vol) twice. Subsequently, 1/10 volume of 3 M sodium acetate and 2 volumes of ice-cold absolute ethanol were added with an overnight incubation at −20°C to precipitate DNA. The precipitated DNA was purified by the addition of 1 mL 70% ethanol. After the evaporation of the ethanol, TE buffer (50 μL) was used to resuspend DNA. The size of the DNA was determined with 0.5 % agarose gel. DNA concentrations were determined by the Nanodrop 2000c spectrophotometer.

### Whole genome sequencing and de novo assembly

Approximately 500 ng high-molecular-weight (HMW) DNA was sent to Roy J. Carver Biotechnology Center, DNA Services Laboratory (University of Illinois at Urbana-Champaign, Urbana, Illinois, United States) for whole genome sequencing. The HMW DNA was sheared with a Megaruptor 3 to an average fragment length of 13 kb. Sheared DNA was converted to a library with the SMRTBell Express Template Prep kit 3.0 from PacBio. The library was sequenced on a shared SMRTcell 8M on a PacBio Sequel IIe using the CCS sequencing mode and a 30-hour movie time. CCS analysis was done in an instrument with SMRTLink V11.0 (PacBio) using the following parameters: ccs --min-passes 3 --min-rq 0.99.

Read quality was evaluated using FastQC v0.11.8^45^. SeqKit v2.0.0^46^ was used to calculate the statistics of the sequencing files for each microbial isolate. Read number, sum of the read length, minimum read length, average read length and maximum read length were shown in **Supplementary Table 4.** Flye v2.9^55^ was used to assembly the reads (enough for 50 fold coverage) chosen using the parameters: --asm-coverage. Assembly quality and completeness were evaluated using QUAST v5.0.2^56^ and BUSCO v5.5.0^57^ respectively. Annotations were performed using Prokka v 1.14.6^58^. CGView Server was used to make the circular genome maps^59^. The 16S rRNA genes were extracted by SnapGene v6.2.1 and used to determine the species by doing the BLAST in NCBI.

### Comparative genomics

Average nucleotide identity (ANI) was calculated using pyani v0.2.10^60^ with the third-party tool MUMmer4^61^. Roary v3.13.0^62^ was used to generate a core gene alignment. The alignment was done via the parameters: -e –mafft. The best model for phylogenomics was calculated by using ModelTest-NG^63^. RAxML-NG^64^ was used to infer the bootstrapping tree with the parameter: --bs-trees autoMRE. An online tool (iTOL) was used for the tree display and annotation^65^. The web application pyGenomeViz v0.4.0^66^ was used to visualize the nucleotide sequence similarity with the GenBank format files as inputs based on MUMmer4. A Venn diagram was generated using *ggvenn* package^67^.

### Metagenomics (MAGs)

Genomes from human gut microbiome datasets were downloaded from the following nine different sources: 32,277 genomes from Zeng et al. 2022^68^, 1,200 genomes from Wilkinson et al. 2020^69^, 120 genomes classified as *C. scindens* from Almeida et al. 2020^70^, 1,381 genomes from Tamburini et al. 2021^71^, 154,723 genomes from Pasolli et al. 2019^72^, 4,997 genomes from Merrill et al. 2022^73^, 2,914 genomes from Lemos et al. 2022^74^, 4,497 genomes from Gounot et al. 2022^75^, and 31 genomes from NCBI. The GTDB-Tk (version 2.1.1) classify workflow (classify_wf) was run on all 202,140 genomes, which resulted in the identification of 224 *C. scindens* genomes. Custom HMMs were generated for the proteins DesA, DesB, and DesF by using experimentally verified sequence. Briefly, in order to create the protein alignments needed to generate HMM profiles, muscle (5.1.linux64) was used with default parameters to align all amino acid sequences for each of the proteins followed by the hmmbuild function of the HMMER (version 3.3.2) package to generate the HMM profiles. Trusted HMM cutoffs were generated for each of the proteins based on the maximum F-scores based on searches with orthologous proteins. The 224 identified *C. scindens* genomes were translated into amino acid sequences using Prodigal (version 2.6.3). The generated HMM profiles were queried against the amino acid sequences for the 224 genomes using hmmsearch of the HMMER package with the flag --cut_tc.

### Phylogenetics (DesF and DesG)

The DesF protein sequence of *C. scindens* (accession number WP_220430766.1) and the DesG protein sequence of *P. lymphophilum* API-1 (this work) were used as queries for BLASTP searches against the NCBI’s NR protein database. The protein sequences from the top one thousand (DesF) or five hundred (DesG) results were aligned using MUSCLE v. 5.1 with default parameters^76^. Ambiguously aligned positions were removed with Gblocks v. 0.91b^77^, allowing up to 50% of the sequences in a column to contain gaps and using minimum length of a conserved block size of 5. Each phylogeny was inferred by IQ-TREE v. 2.2.2.6^78^ using two independent runs, one thousand ultrafast bootstrap pseudoreplicates (optimized by NNI on the bootstrap alignment), one thousand SH-like approximate likelihood-ration test replicates, and extended substitution model selection (including Lie Markov models). Trees were edited in TreeGraph2^79^, drawn in Dendroscope v. 3.8.10^80^, and final cosmetic adjustments were performed in Inkscape (www.inkscape.org).

### Computational Structural Biology

#### Protein Modeling

The models for desF and desG protein structures were predicted using AlphaFold version 2.3.2^22^, employing all five available parameter sets to generate five models each, resulting in a total of 25 predictions per sequence. The QwikFold plugin in VMD^23^ was used to set up the computational experiments and post-process the results, with calculations run using the Cybershuttle^81^. QwikFold was also utilized to align the models for visual inspection, addressing per-residue confidence as measured by pLDDT. The predicted aligned error (PAE) matrices were inspected to assess confidence in the predicted structures. Additionally, both structures were solved using a homology modeling strategy. The enzyme models were constructed using MODELLER^82^, which employs spatial restriction techniques based on 3D-template structures. The best model was selected by analyzing stereochemical quality using PROCHECK^83^ and overall quality using the ERRAT server^84^. VMD was then used to compare the structural models, and all predicted structures were used for the docking strategy.

#### Docking

Using BLAST^85^, we obtained homologous structures (PDB IDs: 4ILK, 4EJ6, 4A2C, 3QE3, 3GFB, 2DQ4, 2DFV, 2D8A, 1PL7, 1E3J) from the protein data bank (PDB). The design of the ligands (NADP+, testosterone, and epiT) was carried out using Molefacture, the small molecule design suite in VMD^86^. The alignment and placement of both NADP+ and the steroid molecules (epiT in desF and testosterone in desG) in their binding sites were performed using VMD. Employing advanced run options in QwikMD^24^, the structures of the ligands were minimized in the pockets along with nearby enzyme residues, while maintaining the structure of most of the enzyme as static. PyContact^87^ was then used to analyze the contact interface.

#### Molecular Dynamics Simulations

MD simulations were performed using the GPU-accelerated NAMD^25^ molecular dynamics package. The simulations assumed periodic boundary conditions in the NpT ensemble, with temperature maintained at 300 K using Langevin dynamics for temperature and pressure coupling, the latter kept at 1 bar. A distance cut-off of 12.0 Å was applied to short-range non-bonded interactions, while long-range electrostatic interactions were treated using the particle-mesh Ewald (PME) method^88^. The equations of motion were integrated using the r-RESPA multiple time step scheme to update the van der Waals interactions every step and electrostatic interactions every two steps. The integration time step was set to 2 fs. Before MD simulations, the system underwent energy minimization and a short MD run protocol for 50,000 steps (1,000 steps of minimization followed by 1,000 steps of MD, repeated 25 times). An MD simulation with position restraints on the protein backbone atoms 7 Å or more away from the ligands was performed for 10 ns. To allow for total system relaxation and ensure ligand stability in the pockets, a 100 ns equilibrium simulation without external forces was performed.

#### MD Analysis

Analyses of MD trajectories were carried out using VMD and its plugins^23^. Surface contact areas of interacting residues were calculated using PyContact^87^. The generalized dynamical network analysis tool was employed to perform dynamical network analysis^89^.

### Statistical analysis

Statistical analyses were performed with R version 4.3.0^90^. Data are shown as means ± standard deviations (SD) when data are normalized. Data are shown in median and interquartile ranges when skewed. Categorical data are shown as counts and percentages. Differences between groups were analyzed by t-test when data are normal or Mann-Whitney U test when not normally distributed. Differences in categorical data were analyzed using chi-square test. The *P* values for multiple tests were corrected using Benjamini-Hochberg false-discovery rate (FDR). A *P* value ≤ 0.05 was considered statistically significant.

## Data and materials availability

The raw RNA-Seq reads are available at the National Center for Biotechnology Information (NCBI) with accession numbers PRJNA1108660 (Experiment one, *desF*), PRJNA1108687 (Experiment two, *desF*), and PRJNA1115127 (*desG*) respectively. The raw genome sequencing data are available at the NCBI with accession numbers PRJNA1108738 (*P. lymphophilum* strains) and PRJNA1026650 (*C. scindens* strains) respectively.

All scripts and HMMs used in the metagenomic analyses are available at https://github.com/AnantharamanLab/Clostridium_scindens_mining.

## Acknowledgements

J.M.R. would like to express gratitude to both the Cancer Center at Illinois and the Center for Advanced Study at Illinois for the financial support and protected time to pursue this work. We would like to thank Alvaro Hernandez, Director of DNA Services Facility at the Roy J. Carver Biotechnology Center at Illinois and Christopher J. Fields, Director of High-Performance Biology Computing at the Roy J. Carver Biotechnology Center for nucleic acid sequencing and bioinformatic assistance. We thank Kristen M. Flatt for SEM images of bacterial isolates. We thank Elizabeth Tang for collecting urine samples. We thank Peter M. Yau for Proteomic Analysis. Figures were created with BioRender.com. All chemical structures were created with Chemdoodle.

## Funding

This work was supported by grants from the National Institutes of Health (R01 GM145920-01 [J.M.R., I.C., H.R.G.], R01 GM134423-01A1 [J.M.R.], R01 CA287126 [K.S.S., J.M.R., J.I., I.C.] R03 AI147127-01A1 [J.M.R. J.M.P.A,]), Cancer Center at Illinois Seed Grant [J.M.R, J.I., J.E. Jr, H.R.G.], Prostate Cancer Foundation grants 16CHAL13 [K.S.S.] and 23CHAL13 [K.S.S., J.M.R., J.I., I.C.], Department of Defense Prostate Cancer Research Program grant W81XWH-20-1-0274 [K.S.S.], as well as UIUC Department of Animal Sciences Matchstick grant, and Hatch ILLU-538-916. F.F. was supported through a Fulbright Fellowship. B.B. was supported by NIGMS Diversity Supplement on R01 GM134423. J.M.R. was supported by an Associateship with the Center for Advanced Study at Illinois during the study period. Computational work supported by the National Science Foundation grant MCB-2143787 and NIH R24 GM145965 [R.C.B.].

## Author Contributions

J.M.R. conceptualized and supervised the overall study and wrote the first draft and edited subsequent drafts. J.I., and K.S.S. supervised cell culture and clinical aspects of the study, respectively, and contributed to the second and subsequent drafts. J.W.E Jr., H.R.G., P.B.H, and I.C. provided guidance and supervision of key aspects of the study. J.M.R. prepared bacterial RNA for RNA-Seq. T.W. performed comparative genomic analysis, transcriptome analysis, heterologous expression, and LC/MS analysis. S.A. performed cell culture experiments, urine culturomics and LC/MS analysis. J.M.R., J.W.L. and B.B. performed cloning and heterologous expression experiments. Y.J. developed co-culture bead methodology. B.B. performed characterization of DesG and orthologs. Y.O.C. and J.M.P.A. performed comparative genomic analysis and phylogenetic analysis. F.F. performed bioinformatic analysis. G.Y. recruited patients. P.M., D.D., collected patient urine samples. A.M.B. and K.A. performed metagenome analysis. A.C and S.E.E performed qPCR quantification of *desF* in clinical samples. S.H., J.D.K and S.I. prepared samples for or performed NMR analysis. R.C.B. performed and analyzed the computational structural biology work. J.M.R., K.S.S., J.I., T.W., S.A., and S.L.D. analyzed data and wrote the manuscript with input from all the co-authors.

## Competing Interests

The authors declare no competing interests.

## Correspondence and requests

Jason M. Ridlon; Joseph Irudayaraj; Karen S. Sfanos;

## Extended Data Figures

**Extended Data Figure 1.**
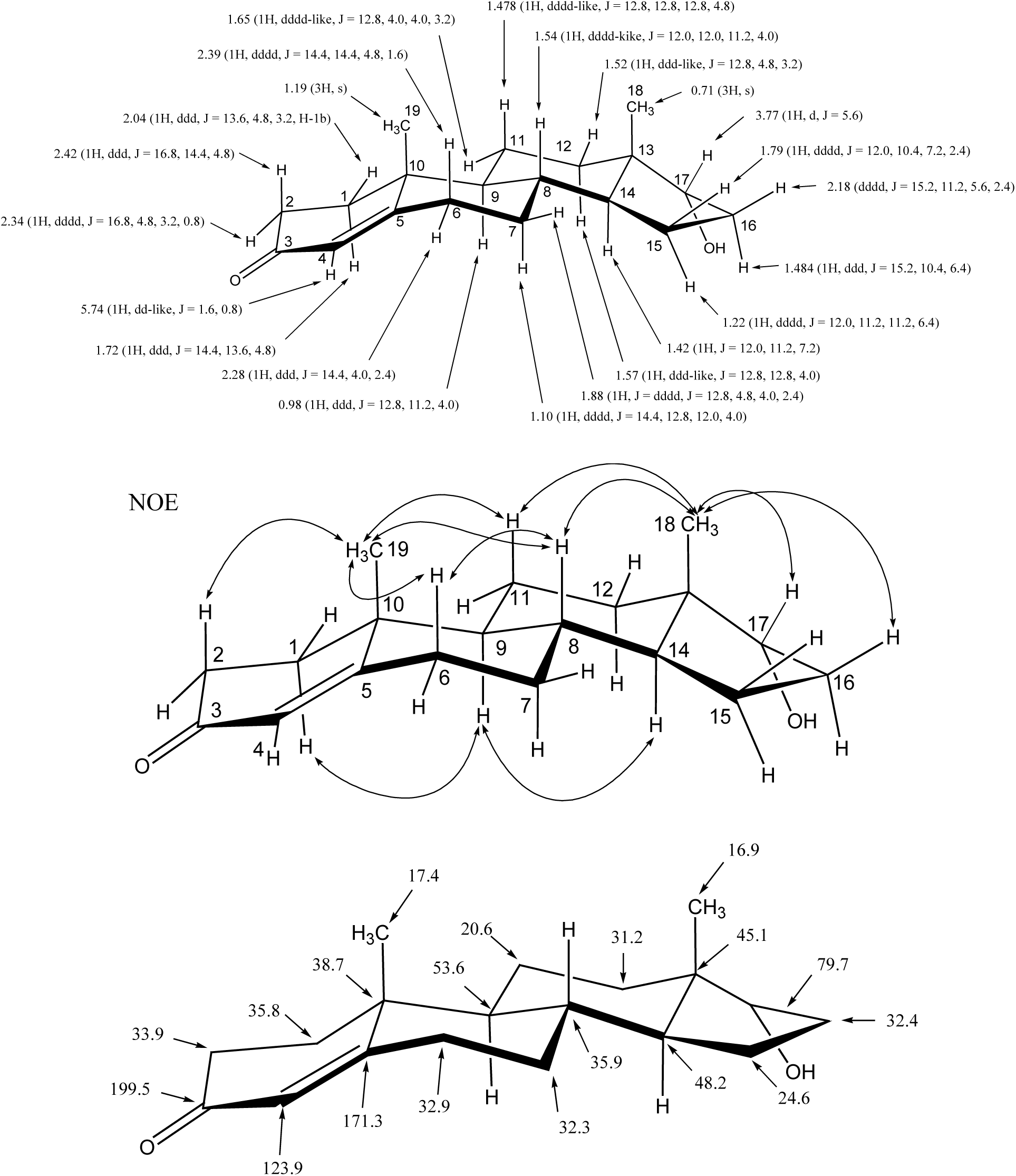
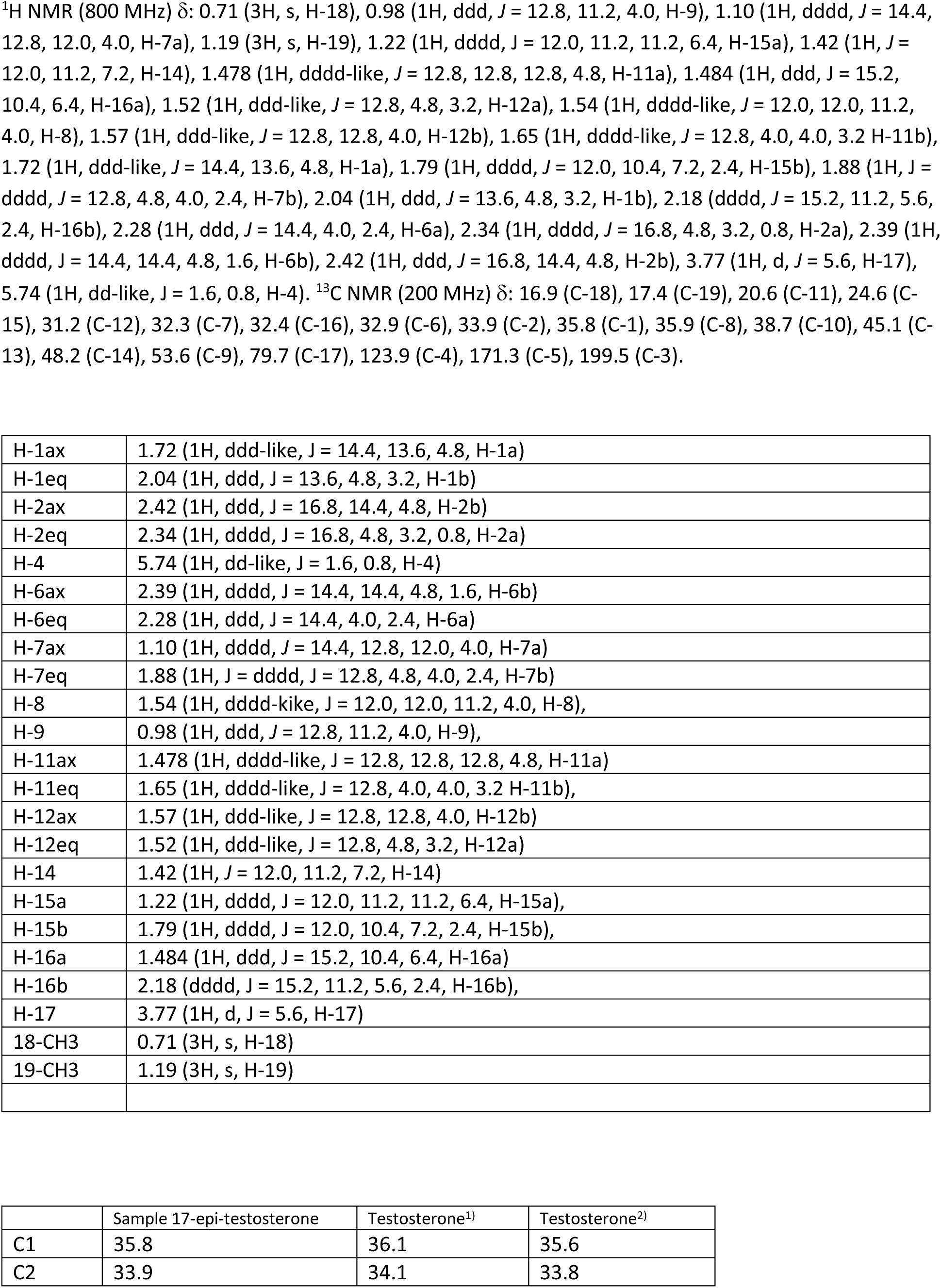
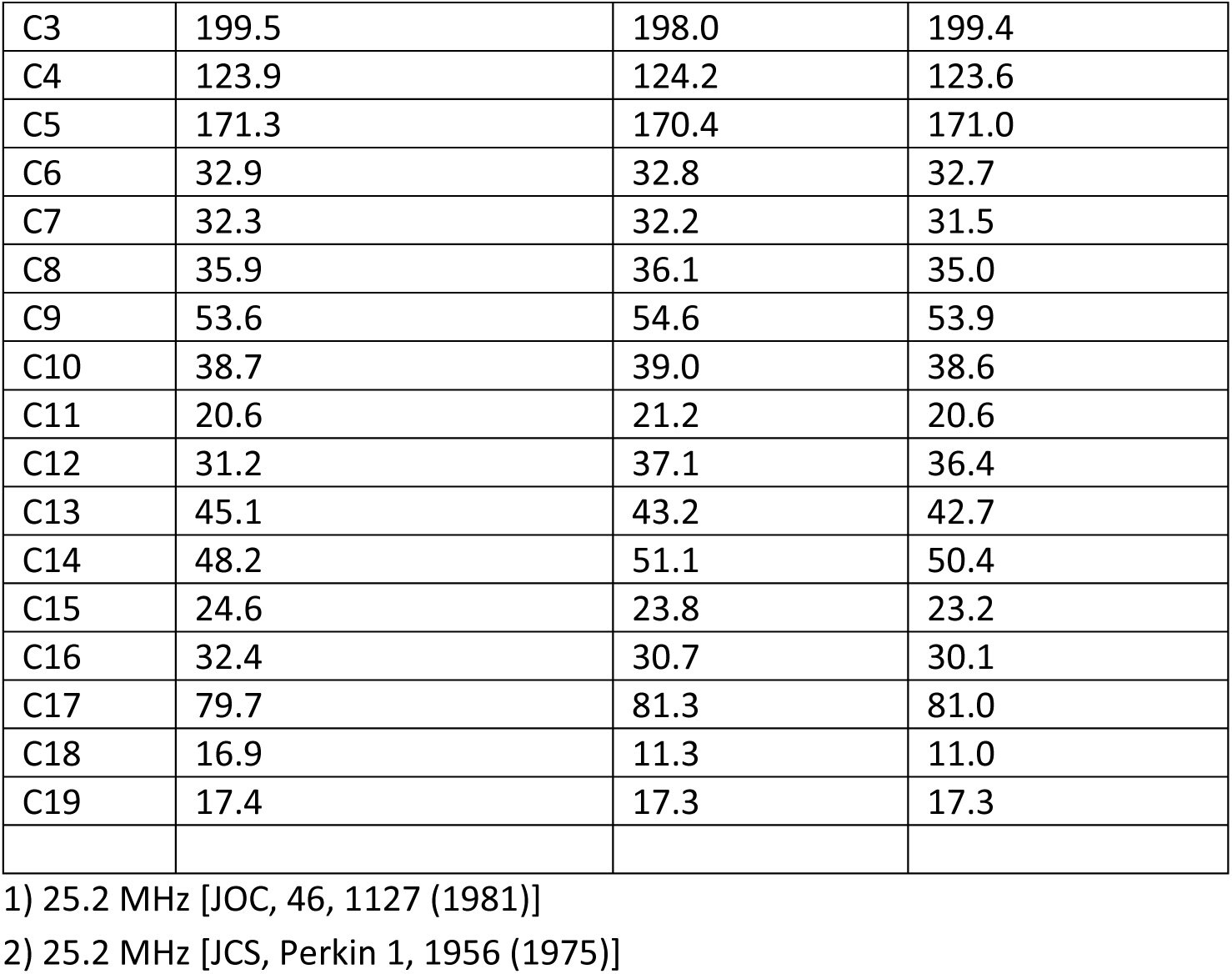
Proton and Carbon NMR analysis of purified reaction product of androstenedione in cultures of *Clostridium scindens* VPI12708. ^1^H and ^13^C-NMR spectroscopic data were obtained on a JNM-ECA800 (JEOL, Ltd., Tokyo, Japan) instrument operated at 800 and 200MHz, respectively, with CDCl_3_ as the NMR solvent. Chemical shifts were expressed in d (ppm), and coupling constants *J*_H,H_ are given in Hz. ^1^H–^1^H nuclear overhauser effect spectroscopy (NOESY), ^1^H-^1^H correlation spectroscopy (COSY), ^1^H-^13^C heteronuclear single-quantum correlation spectroscopy (HSQC), and ^1^H-^13^C heteronuclear multiple-bond correlation spectroscopy (HMBC) spectra were obtained using gradient-selected pulse sequences. The ^13^C distortionless enhancement by polarization transfer (135°, 90°, and 45°) spectra were measured between CH_3_, CH_2_, CH, and coherence based on their proton environments.

**Extended Data Figure 2.**
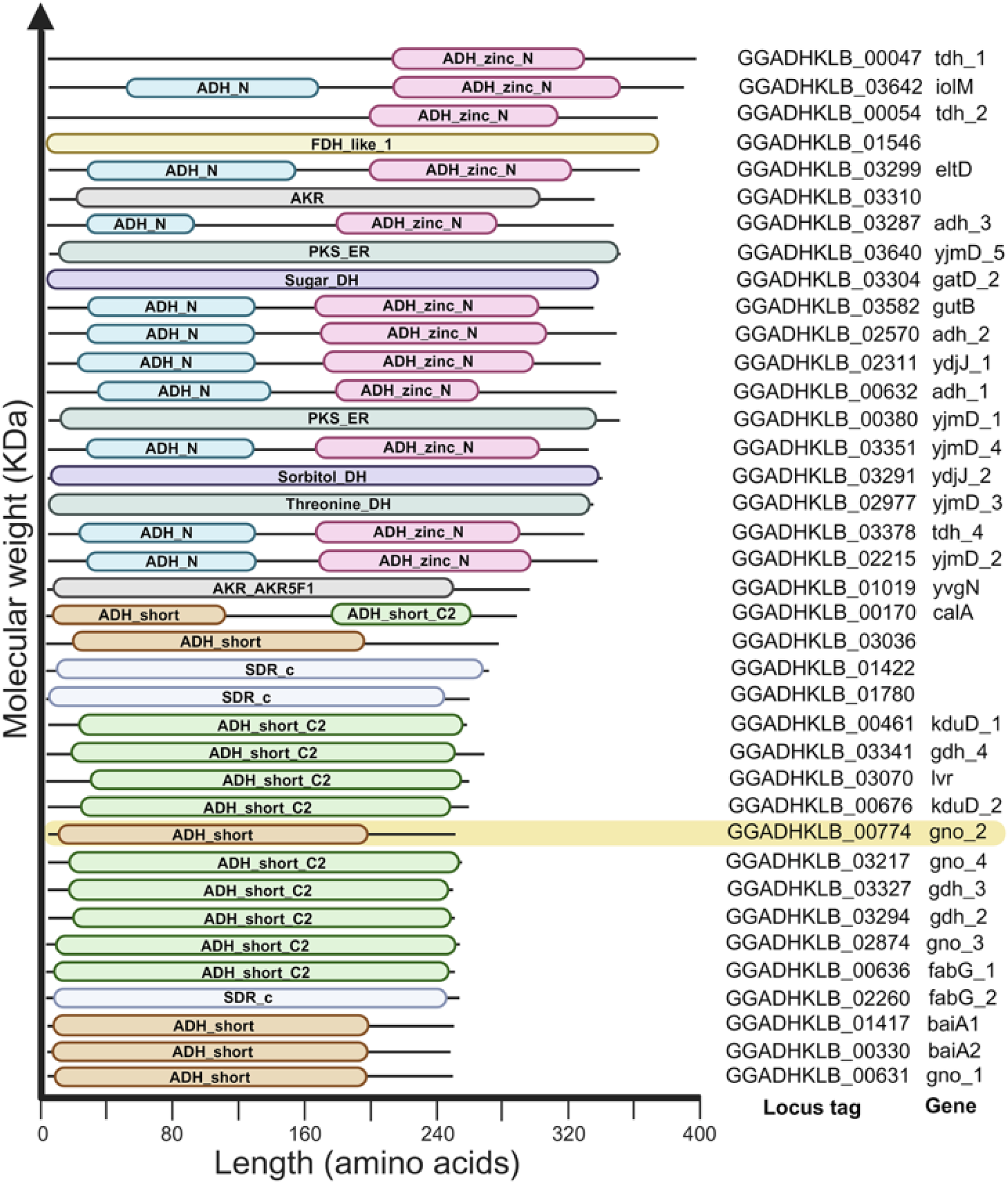
Comparative genome analysis of *Csci*35704 and *Csci*12708 revealed reductase candidates for 17α-HSDH. Highlighted in yellow is the candidate significantly upregulated in RNA-Seq dataset (See main text **Figure 1**).

**Extended Data Figure 3.**
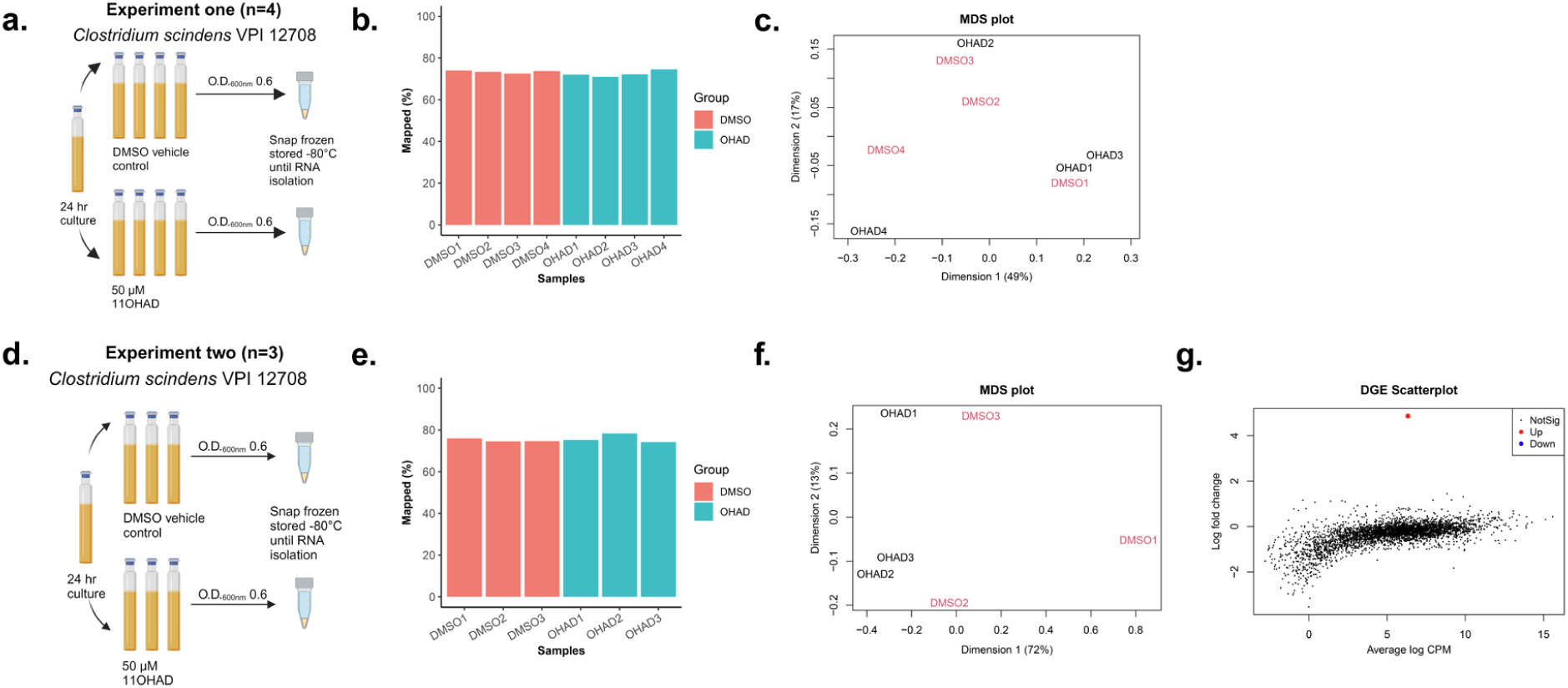
RNA-seq analysis to identify the gene encoding 17ɑ-hydroxysteroid dehydrogenase involved in epitestosterone formation by *Clostridium scindens*. **a.** *In vitro* induction of the gene encoding 17ɑ-hydroxysteroid dehydrogenase by addition of 50 µM 11OHAD. DMSO was used as the vehicle control (Experiment one, n=4). **b.** Percentages of the reads mapped to the reference genome of *Clostridium scindens* VPI 12708 (Experiment one). **c.** MDS plot showed the gene composition by *in vitro* induction of Experiment one. **d.** *In vitro* induction of the gene encoding 17ɑ-hydroxysteroid dehydrogenase by addition of 50 µM 11OHAD (Experiment two, validation experiment, n=3). **e.** Percentages of the reads (Experiment two) mapped to the reference genome of *Clostridium scindens* VPI 12708. **f.** MDS plot showed the gene composition by *in vitro* induction (Experiment two). **g.** Differential gene expression scatter plot of Experiment two. Significantly upregulated genes in red, downregulated genes in blue, and not differentially regulated in black.

**Extended Data Figure 4.**
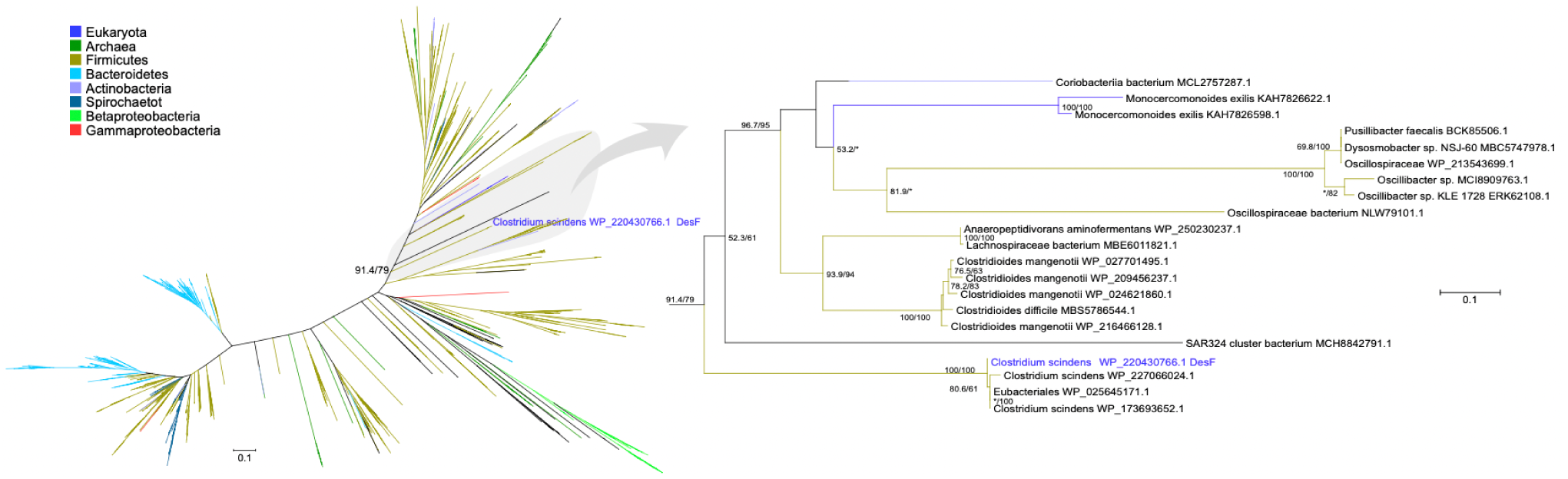
Maximum likelihood phylogeny of DesF. Tree was colored by taxonomic affiliation, according to the included key. Radial tree shows the complete one-thousand sequence tree, and the region near *C. scindens* is shown as a phylogram. Numbers near nodes are branch support values (SH-like approximate likelihood-ration test and ultrafast bootstrap), with only values greater than 50 shown (an * denotes that one of the values is under 50 but the other is not).

**Extended Data Figure 5.**
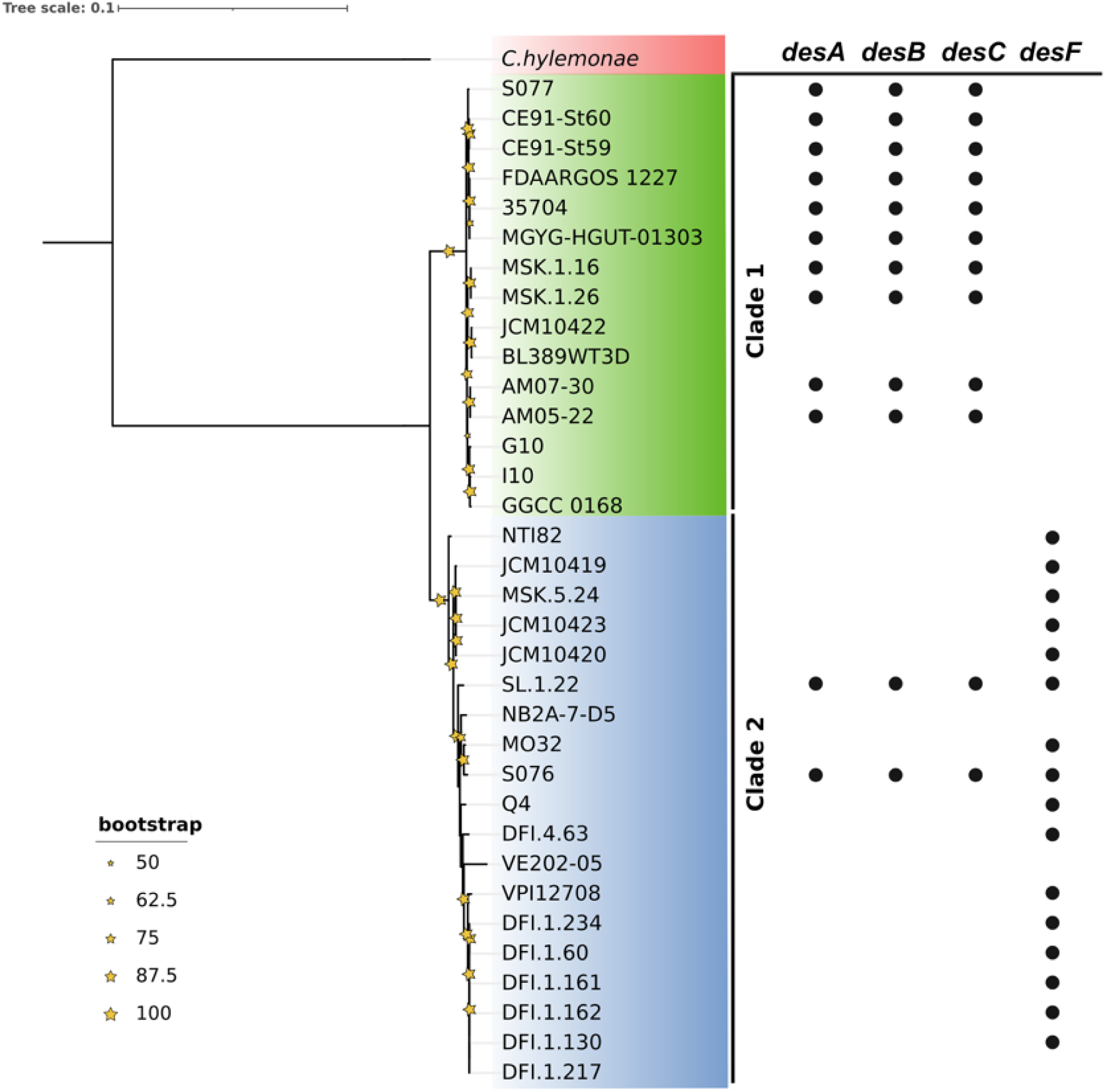
Phylogenomic and diversity of *des* gene presence in strains of *C. scindens*. The formation of two clades is shown, Clade 1 (green) includes 15 strains and Clade 2 (blue) 19 strains. Bootstrap support values above 50% are shown in yellow stars at nodes. Dots show *des* gene presence in strains of each clade.

**Extended Data Figure 6.**
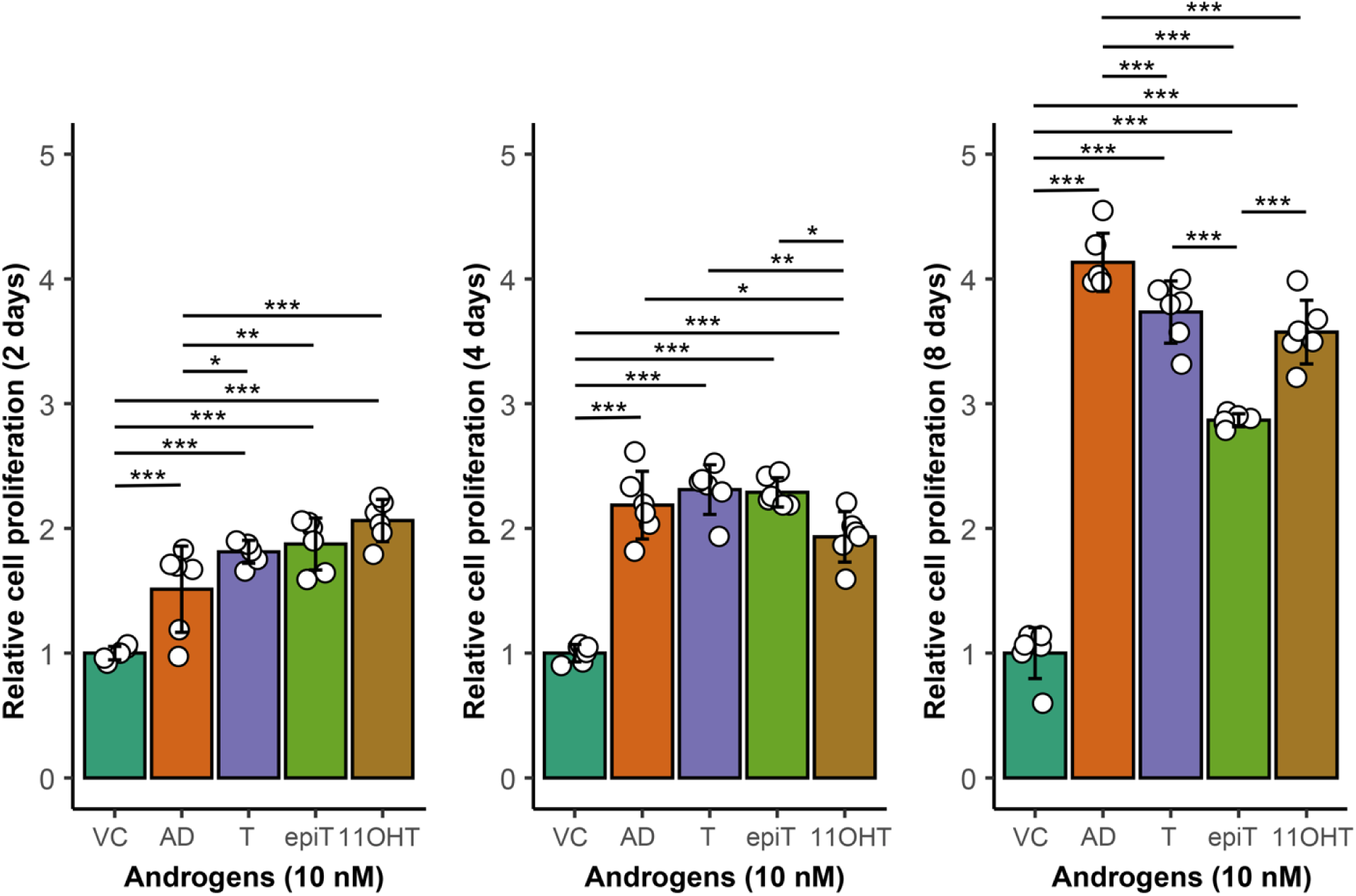
Steroid-dependent proliferation of VCaP cell line. Data are shown with mean ± standard deviation (n=6). *P* values were calculated by unpaired *t*-test and Benjamini-Hochberg correction, * *P <* 0.05, ** *P <* 0.01,*** *P* < 0.001. VC: Vehicle control (0.5% methanol v/v); AD: Androstenedione; T: Testosterone; epiT: Epi-testosterone; 11OHT: 11β-hydroxy-testosterone.

**Extended Data Figure 7.**
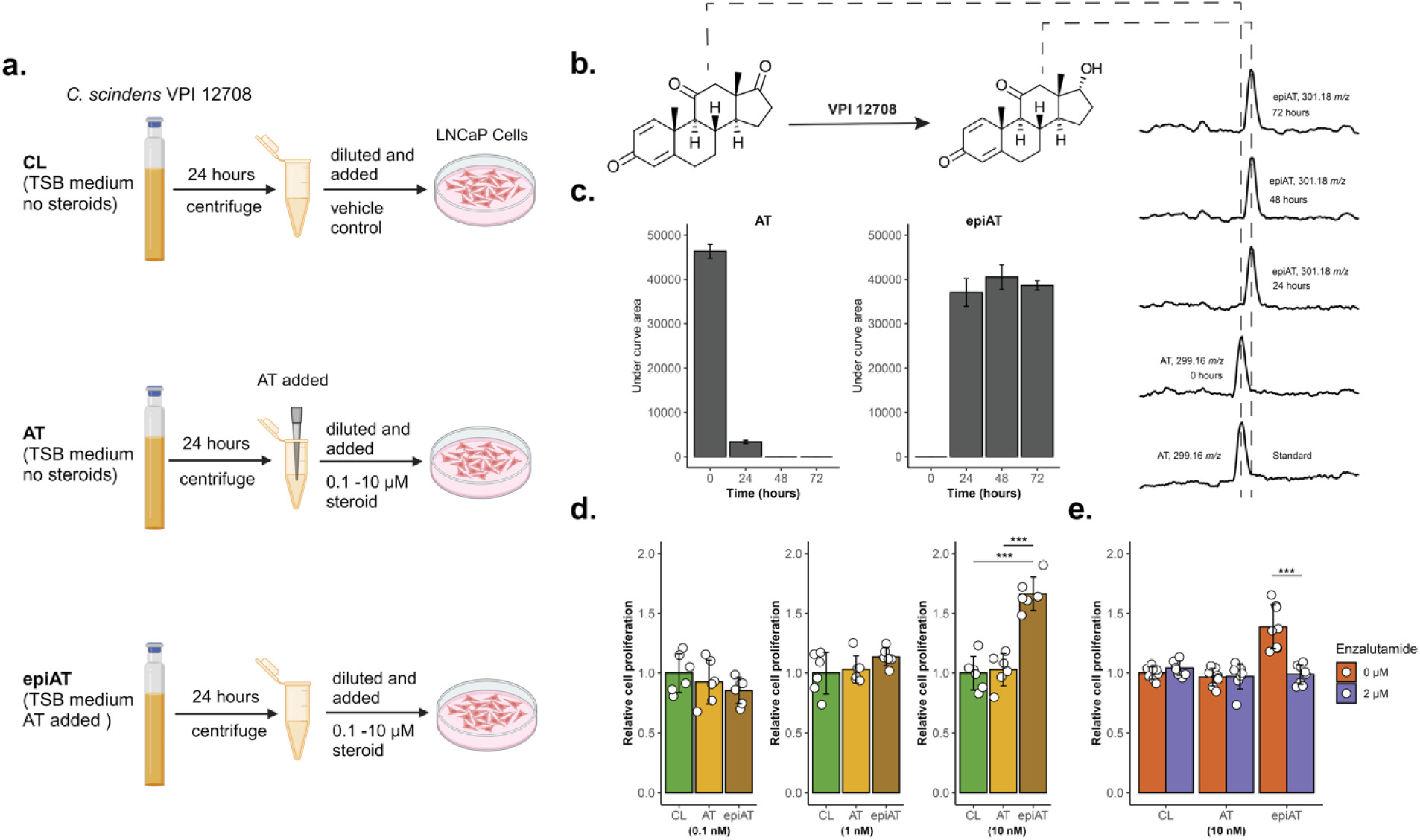
*Clostridium scindens* generates androgenic derivative of epitestosterone from 1,4-androstadiene-3,11,17-trione (AT). **a.** *In vitro* incubation of *Clostridium scindens* VPI 12708 with 50 µM AT as substrate. The medium culture after removing the bacterial cells was diluted and used for the LNCaP cell line promotion effect study. **b.** A proposed biochemical pathway by which *Csci*12708 convert AT to epiAT. **c.** LC/MS chromatographs of AT and epiAT at time 0, 24, 48 and 72 h and the change of the under-curve area of AT and epiAT overtime (n=3). **d.** Dose-dependent proliferation of LNCaP cells in the presence of medium culture control (CL), medium culture control with AT (AT), and metabolite of AT in the medium culture (epiAT) at 0.1 nM, 1.0 nM and 10.0 nM (n = 6). **e.** Enzalutamide effect on the LNCaP cell line proliferation in the presence of medium culture control (CL), medium culture control with AT (AT), and metabolite of AT in the medium culture (epiAT) at 10.0 nM (n = 8). Data are shown with mean ± standard deviation. *P* values were calculated by unpaired *t*-test and Benjamini-Hochberg correction, * *P <* 0.05, ** *P <* 0.01,*** *P* < 0.001.

**Extended Data Figure 8.**
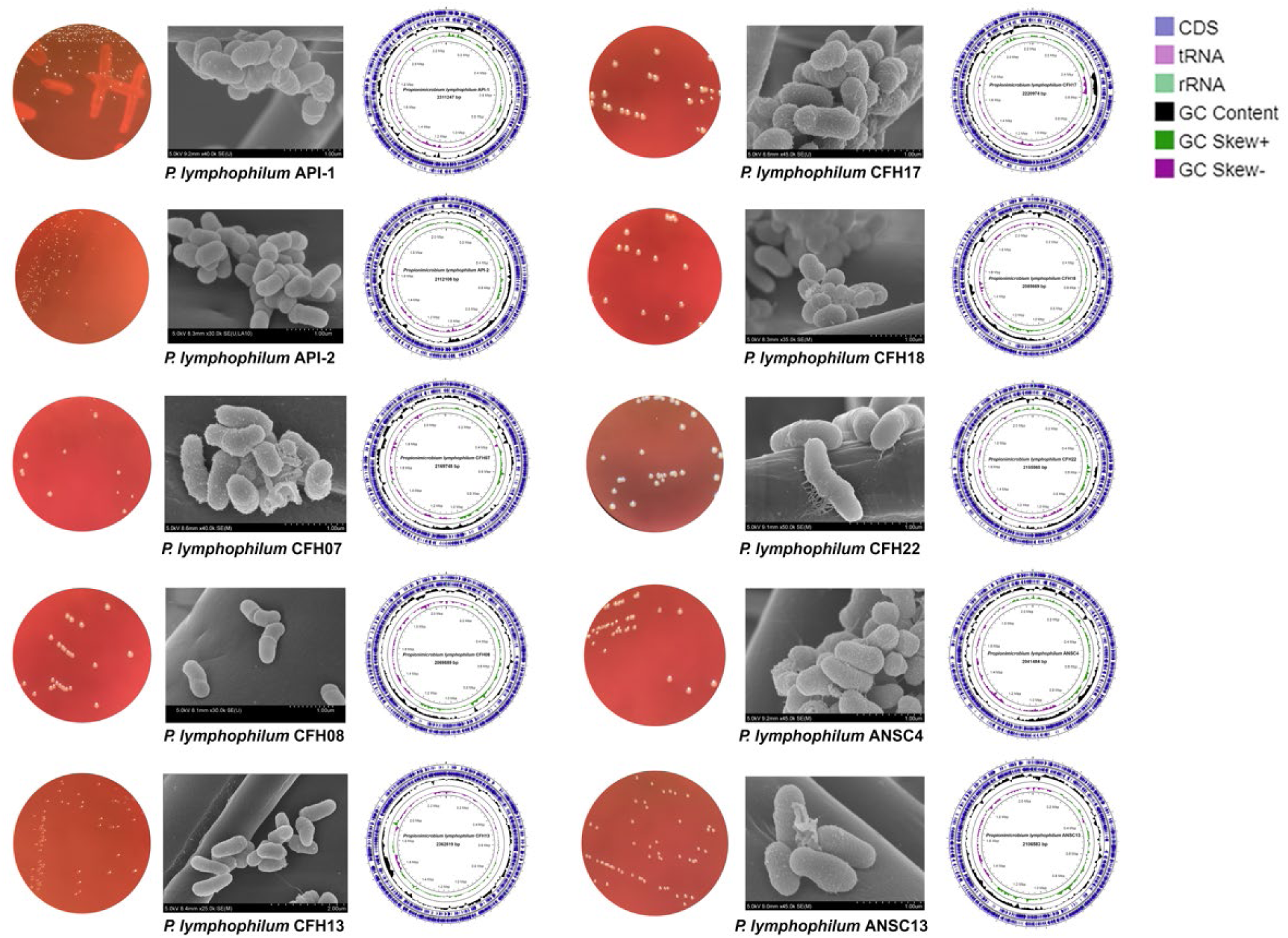
Bacterial colony morphology, SEM images and the circular genome maps of the androgen-producing *Propionimicrobium lymphophilum* strains from men diagnosed with prostate cancer and age-matched control males.

**Extended Data Figure 9.**
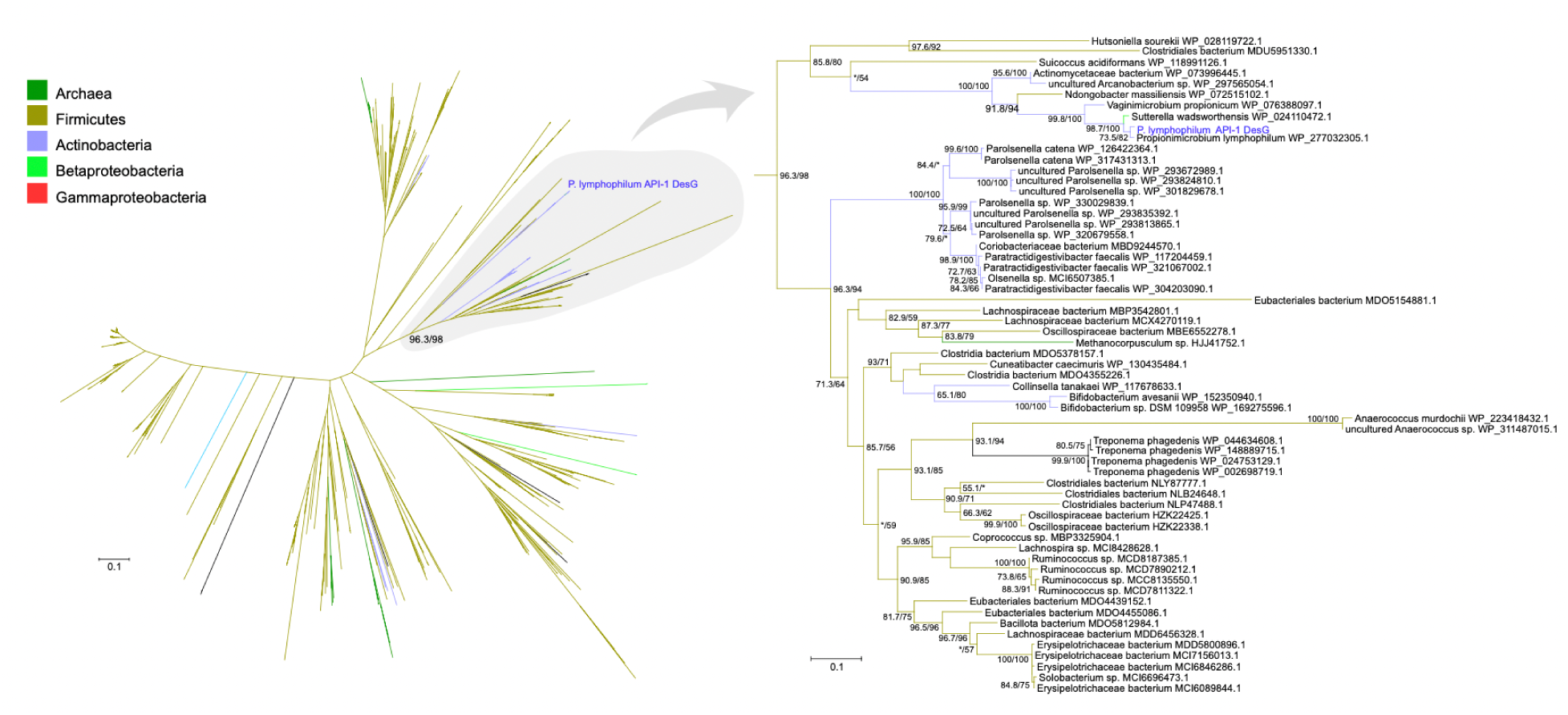
Maximum likelihood phylogenetic analysis of DesG from *Propionimicrobium lymphophilum* strain API-1. Tree was colored by taxonomic affiliation, according to the included key. Radial tree shows the complete five-hundred sequence tree, and the region near *P. lymphophilum* is shown as a phylogram. Numbers near nodes are branch support values (SH-like approximate likelihood-ration test and ultrafast bootstrap), with only values greater than 50 shown (an * denotes that one of the values is under 50 but the other is not).

**Extended Data Figure 10.**
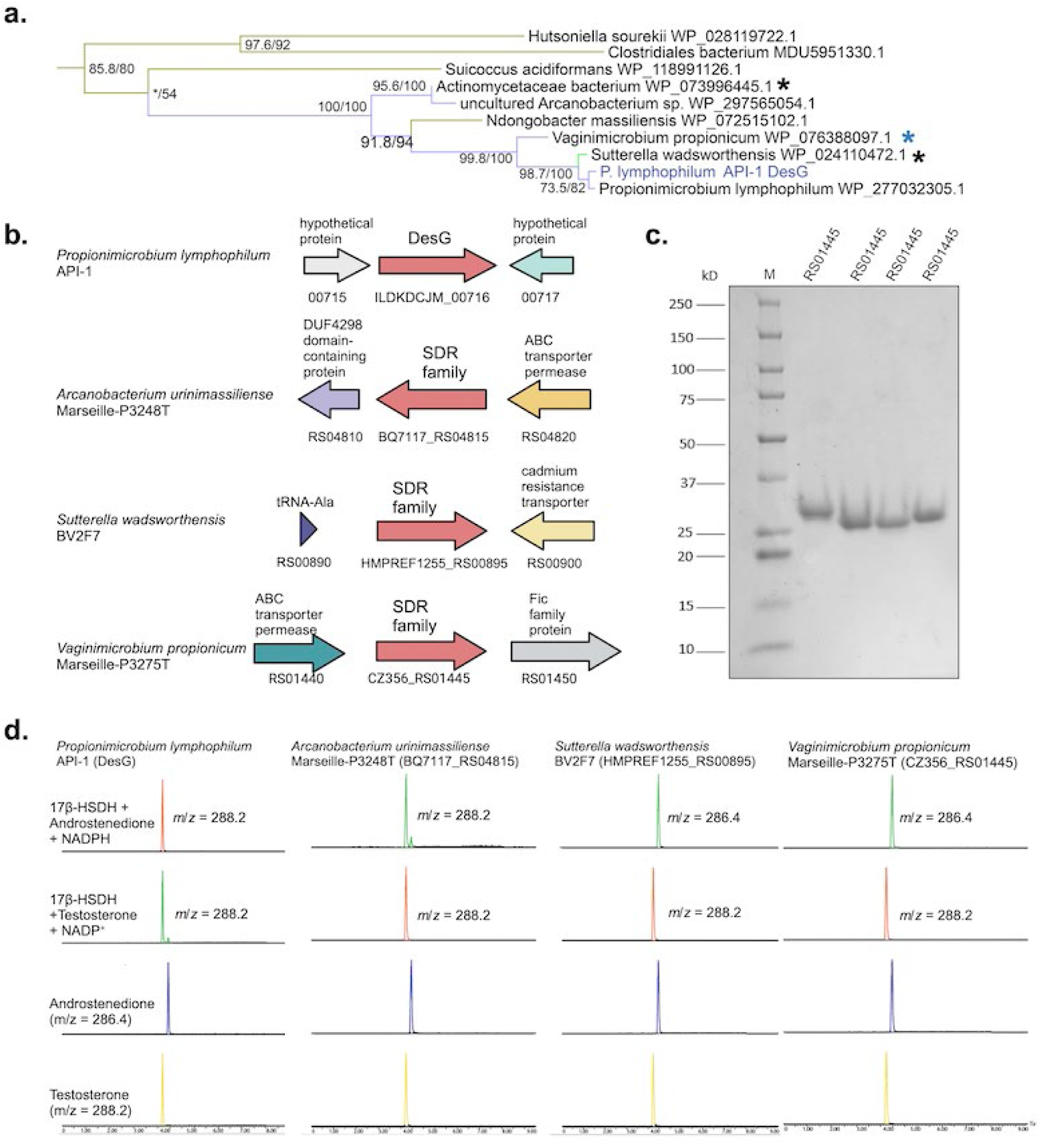
Functional sampling of *desG* homologs for 17β-HSDH activity. **a.** Zoom in of DesG branch with proteins chosen for functional characterization (asterisks). **b.** Genomic context of *desG* candidates from taxa represented in DesG branch. **c.** SDS-PAGE of affinity (Streptactin) purified recombinant DesG candidates. **D.** LC/MS traces after enzyme assay containing either DesG candidate (10 nM) + NADPH (200 µM) and androstenedione (50 µM), or DesG candidate (10 nM) + NADP+ (200 µM) and testosterone (50 µM) incubated for 6 h before steroid extraction. Authentic androstenedione and testosterone standards were also included.

**Extended Data Figure 11.**
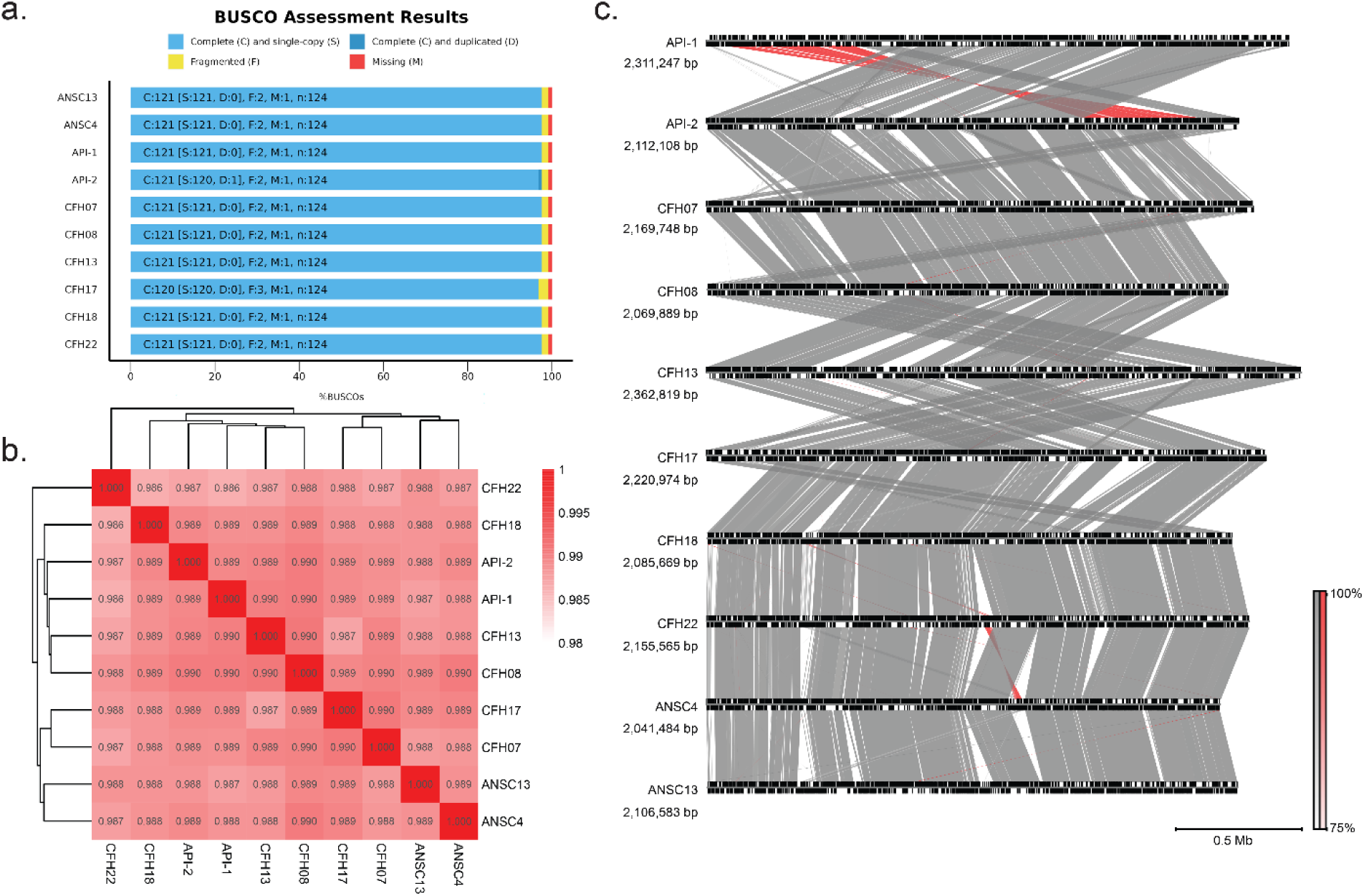
Genome quality metrics (a.), average nucleotide identity, (b.) and synteny between *Propionimicrobium lymphophilum* strains (c.) isolated from human male urine.

**Extended Data Figure 12.**
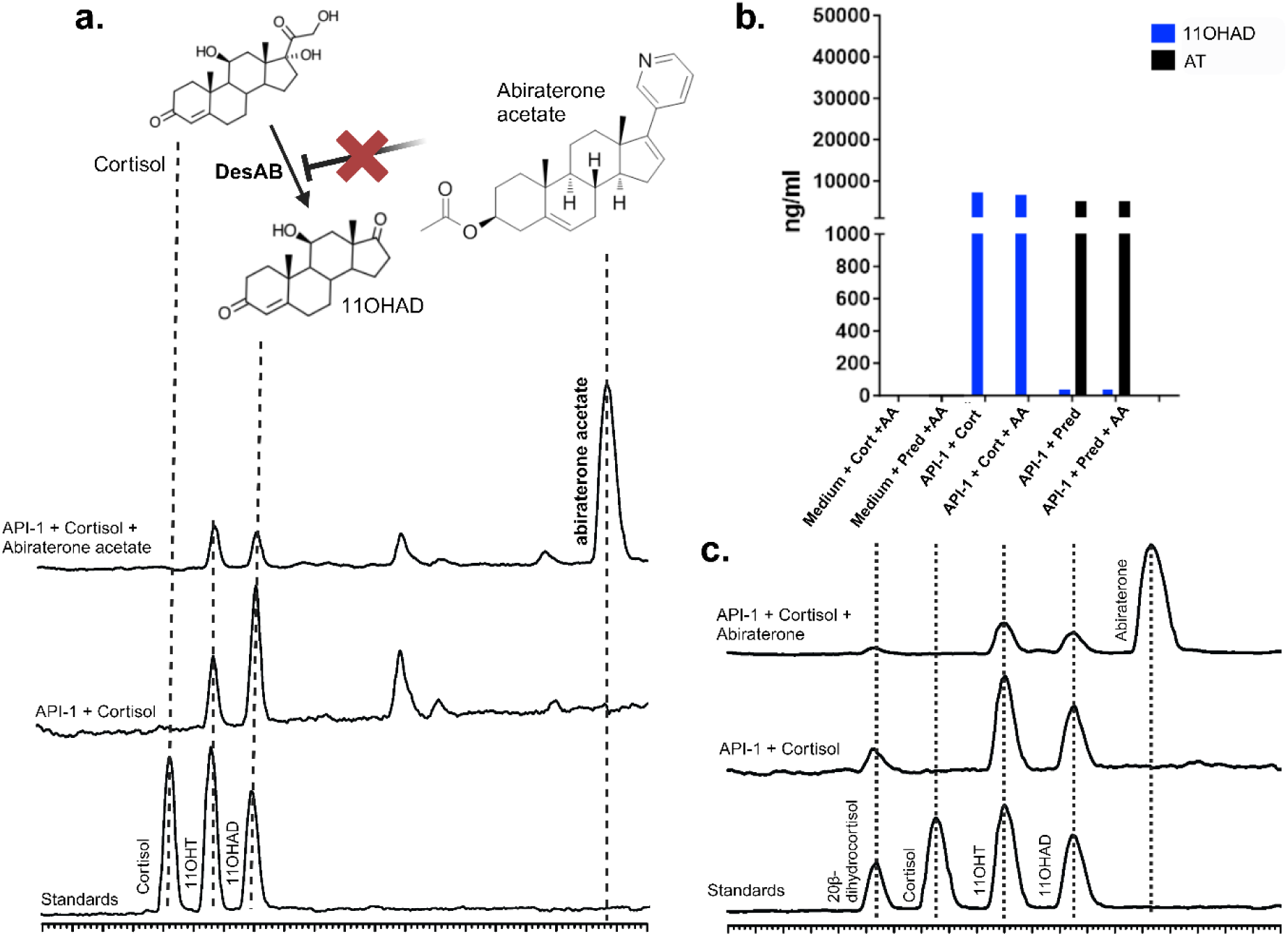
Abiraterone (A) and abiraterone acetate (AA) are not inhibitors of bacterial steroid-17,20-desmolase (*desAB*). **a.** *P. lymphophilium* API-1 treated with 50 µM AA and then incubated 50 µM cortisol for 48 hours. **b.** API-1 strain treated with AA (1 µM and cortisol or prednisone (50 µM). (C) API-1 treated with 50 µM A and then incubated with 50 µM cortisol for 48 hours.

**Extended Data Figure 13.**
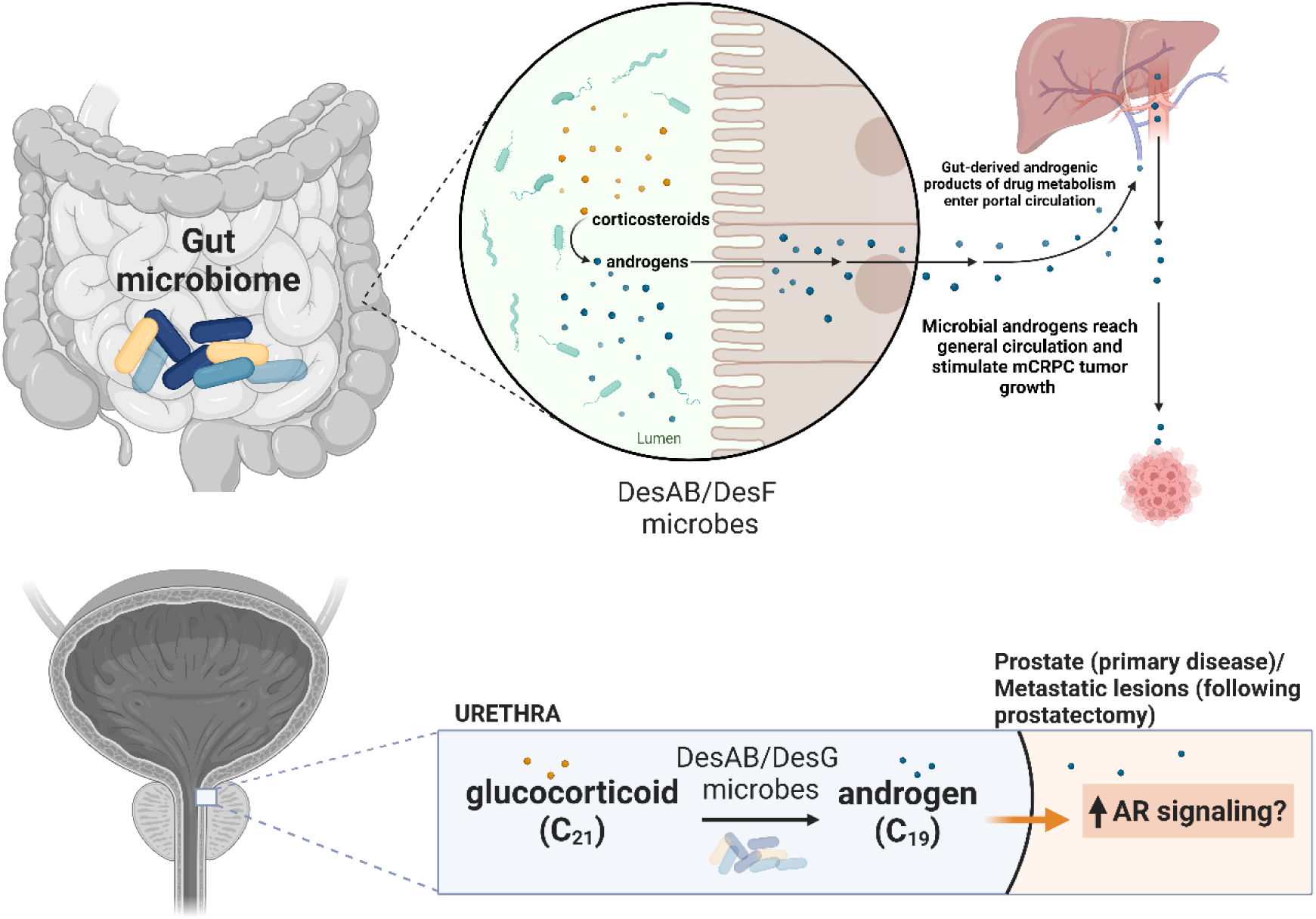
Schematic representation of potential host-microbiome interactions relating to the conversion of glucocorticoids to androgens.

## Notes

### Competing Interest Statement

The authors have declared no competing interest.

